# L-selectin shedding regulates functional recovery and neutrophil clearance following spinal cord injury in a sex-dependent manner

**DOI:** 10.1101/2025.05.28.656699

**Authors:** Miranda E. Leal-Garcia, Mia R. Pacheco, Shelby K. Reid, Victoria S. Tseng, Megan Kirchhoff, Sai Tadiboyina, David Min, Dylan A. McCreedy

## Abstract

During the acute phase of spinal cord injury (SCI), neutrophils infiltrate in large numbers and can exacerbate inflammation, secondary tissue damage, and neurological deficits. L-selectin is a signaling and adhesion receptor that has been shown to facilitate neutrophil recruitment and secondary injury after SCI. During neutrophil activation, L-selectin is typically cleaved or shed from the cell surface and augmenting L-selectin shedding can improve hindlimb recovery and tissue sparing following SCI in male mice. However, it is unclear how endogenous L-selectin shedding regulates neutrophil responses and functional recovery after SCI, particularly when also considering sex as a biological variable. In this study, we investigated the sex-dependent role of endogenous L-selectin shedding in neutrophil function and long-term outcomes in a murine thoracic contusion model of SCI. We found that endogenous L-selectin shedding improves long-term functional recovery and white matter sparing in female, but not male, mice. In addition, we demonstrate that L-selectin shedding alters neutrophil accumulation in a sex-dependent manner. While L-selectin shedding does not mediate neutrophil activation or effector functions, we found that neutrophil clearance is facilitated by L-selectin shedding in female mice alone. These results demonstrate that endogenous L-selectin shedding is a critical and sex-dependent mediator of neutrophil accumulation and clearance, as well as long-term functional outcomes, after SCI.

## Introduction

Spinal cord injury (SCI) is a traumatic life event that can result in permanent paralysis and a decrease in life expectancy^1^. In response to the physical trauma to the spinal cord, damage-associated molecular pattern (DAMPs) molecules and chemokines are released leading to the infiltration of peripheral immune cells (leukocytes)^2^. The accumulation of leukocytes in the spinal cord parenchyma can further exacerbate tissue damage, resulting in even greater subsequent inflammation. Attenuating pro-inflammatory leukocyte responses has great potential as a neuroprotective strategy for SCI. However, the complexity of leukocyte responses in the injured spinal cord, as well as sex differences in inflammation^3–6^, have hindered the identification of effective targets to reduce inflammatory damage and improve long-term outcomes.

Neutrophils are the first peripheral leukocyte to arrive at the injured spinal cord in large numbers, with peak infiltration typically occurring within the first day following SCI in rodents^7–11^. Neutrophils, along with other leukocytes, can be recruited from the vasculature to the site of injury through a process known as transendothelial migration (TEM). During TEM, adhesion molecules and receptors on neutrophils, as well as endothelial cells, facilitate the rolling, arrest, and TEM of neutrophils into the spinal cord parenchyma. Once neutrophils enter the injured spinal cord, they become activated and perform a variety of effector functions, including the release of granules, neutrophil extracellular trap (NET) production and superoxide release^12–14^. Afterwards, neutrophils can be cleared from the inflammatory environment via cell death and phagocytosis by macrophages. Alternatively, neutrophils may re-enter circulation through reverse TEM (rTEM) as a way to resolve inflammation^15–18^. When neutrophils persist in injury sites, resolution of inflammation is hindered leading to further tissue damage^16,19–22^. While neutrophils can worsen tissue damage and long-term neurological outcomes^12–14,23–27^, the sex-dependent processes regulating destructive neutrophil activities in the injured spinal cord remain unclear.

L-selectin is a signaling and adhesion receptor broadly expressed on leukocytes, especially neutrophils. L-selectin facilitates neutrophil rolling along the inflamed endothelium prior to TEM and has been shown to regulate neutrophil recruitment to sites of inflammation^28–31^. In addition, L-selectin also contributes to signaling, activation, and the regulation of neutrophil effector functions^28,30,32–39^. A key feature of L-selectin is the ability to undergo proteolytic cleavage at the membrane proximal domain, resulting in the release of the ligand-binding ectodomain in a process known as “shedding”. The shedding of L-selectin occurs in response to various stimuli and serves as a well-known marker for neutrophil activation^32,40–42^. L-selectin shedding has also been shown to regulate neutrophil function and may be involved in neutrophil cell death and rTEM, thereby, contributing to inflammation resolution and neutrophil clearance^39,43–46^. However, the role of L-selectin shedding in neutrophil responses in the context of SCI has yet to be investigated.

Our previous research demonstrated that L-selectin exacerbates inflammation and long-term neurological deficits in male mice after SCI. Furthermore, we found that augmenting L-selectin shedding improves long-term outcomes and diminishes oxidative stress after SCI in male mice^47^. In this study, we investigate the role of endogenous L-selectin shedding in neutrophil responses and long-term outcomes after SCI while also considering sex as a biological variable. We utilized L(E) mice, which have a mutation in the L-selectin cleavage domain, preventing shedding during neutrophil activation^48^. We demonstrate that genetic loss of L-selectin shedding alters neutrophil accumulation and prolongs neutrophil retention in the injured spinal cord in a sex-dependent manner but does not affect neutrophil effector functions. We also found that abrogating L-selectin shedding worsens long-term recovery and tissue sparing in female, but not male, mice after SCI. Collectively, our findings show that L-selectin shedding regulates neutrophil responses and long-term outcomes in a sex-dependent manner after SCI and further demonstrate the need to consider sex as a biological variable in SCI research.

## Methods

### Mice

All studies were performed in accordance with protocols approved by the Institutional Animal Care and Use Committee at Texas A&M University. L(E) mice were generously provided by Dr. Tom Tedder^48^. Wild type (WT) C57Bl/6N mice were obtained from Charles River Laboratories. Homozygous L(E) mice and WT littermates were generated by breeding heterozygous males and females on a C57Bl/6N background. Homozygous L-selectin KO mice were maintained on a C57Bl/6N background. Genotypes were verified by PCR using a common forward primer (GGGGACATCAGAAGACCAAA) and reverse primers for the WT (GTACCACTGGGG CTGAAAGA) and L(E) alleles (AGGAATGAAGAGGGGGTTGT). Adult male and female mice (12 – 20 weeks old) were used in all experiments. Mice were housed in a climate-controlled room on a 12-hour light/dark cycle with food and water ad libitum. Prior to injury, the mice were housed in groups of two to five. Females were housed in groups of two to three after SCI and males were single housed.

### Spinal cord injury

Mice were anesthetized with 2% isoflurane and a laminectomy was performed at thoracic vertebrate level 9 (T9) to expose the spinal cord. A moderate 70 kDyn contusion injury with a one second dwell was performed using an Infinite Horizons Impactor (Precision Systems and Instrumentation). Prior to impact, the spine was stabilized using clamps placed above and below the injury site (T8 and T10).

Afterwards, the wound was sutured, and the mice received a topical administration of bupivacaine at the incision site. The skin incision then was closed with wound clips and the mice received a subcutaneous injection of saline. During and after surgery, mice were on a water-circulating heat pad set at 37°C to maintain body temperature. For 10 days following SCI, the mice received daily subcutaneous injections of saline and antibiotic (enrofloxacin, 2.5 mg/kg). Mice with endpoints ≤ 24 hours, did not receive antibiotics. Bladder expressions were manually performed twice a day until euthanasia. The body weight was measured at 1, 3 and weekly following SCI until euthanasia. Mice that lost significant body weight (>15%) were provided Nutrical to encourage eating.

### Hindlimb locomotor recovery

The Bass Mouse Scale (BMS) was used to determine hindlimb functional recovery of the mice^49^. To examine their hindlimb locomotor recovery, the mice were placed in an open field and observed for 4 minutes by 2 observers blinded to the genotype. Scores were averaged between the observers. The mice were tested at 1, 3, 7 and weekly thereafter until 5 weeks after SCI.

### White matter sparing

At 35 days post-SCI (dpi), the mice were euthanized using 2.5% Avertin (20 μl/g). The mice were then transcardially perfused with 30 ml of ice-cold 1X PBS followed by 30 ml of ice-cold 4% paraformaldehyde in 1X PBS. The spinal columns were dissected and post-fixed in 30 ml of 4% paraformaldehyde overnight at 4°C. The spinal cords were then dissected and cryoprotected in 30% sucrose in 1X PBS for 2 to 4 days. Then 4 mm centered over the injury site was dissected, embedded, and frozen in optimal cutting temperature medium at −80°C. A series of 25 μm transverse sections spanning the lesion site were made using a Leica CM 1860 cryostat.

Two slides (sections spaced ∼ 250 μm apart) were chosen for eriochrome cyanine staining. Slides were thawed at RT for 10 minutes, then placed on a heat block at 37°C for 10 minutes. Tissue sections were cleared in Citrisolv for 5 minutes. The sections were hydrated in decreasing ethanol solutions for 1 minute each (100%, 100%, 95%, 70%, 50%). The sections were then washed in distilled water for 2 minutes. Tissue sections were then stained in 0.2% eriochrome cyanine stain for 10 minutes then washed in distilled water 3 times for 5 minutes each cycle. The sections were immersed in differentiating solution (5.6% ferric chloride) for 3 minutes then washed in distilled water 3 times for 1 minute each cycle. The sections were dehydrated in using an increasing ethanol gradient (70% for 1 minute, 95% for 2 minutes, 100% x 2 for 1 minute each). Tissue sections were then cleared in Citrisolv 2 times for 5 minutes each cycle. The sections were coverslipped with cytoseal and allowed to dry overnight.

After staining, the slides were imaged on a Leica DM 6B microscope with a 10X objective. Using ImageJ, the white matter was traced to obtain the total area of remaining white matter normalized to the area of the section. The section with the least percentage of spared white matter was selected as the lesion epicenter.

### Neutrophil counting

The animals were euthanized at 1, 3, 14 and 35 dpi and transverse sections of the spinal cord were obtained as described in “white matter sparing”. Two slides (sections spaced ∼ 250 μm apart) were chosen for immunofluorescent staining. The sections were permeabilized in 0.3% Triton-X in 1X PBS and then blocked in 5% normal donkey serum (NDS, Lampire Biological) + 0.1% bovine serum albumin (BSA, Sigma Aldrich) + 0.2% Triton-X in 1X PBS. The sections were labeled with rat anti-Ly6G (Clone 1A8, 1:200 dilution, Biolegend), rabbit anti-NeuN (1:1000 dilution, Millipore) and Fluoromyelin Green (1:300, Invitrogen) overnight at 4°C. The sections were then washed 3 times for 15 min in 1X PBS and stained with AlexaFluor 647 and AlexaFluor 555 conjugated secondary antibodies (1:500, Invitrogen). After staining, the slides were coverslipped and imaged on a Leica DM6B microscope or a Nikon Eclipse upright microscope with a 10X objective. The number of neutrophils were quantified using a semi-automated neutrophil quantification Matlab code^23^ by manually drawing around regions of interest for the entire section, the meninges, the grey matter, and the white matter. The section that contained the highest number of neutrophils was determined to be the lesion epicenter.

### Macrophage accumulation and reactive gliosis

Transverse spinal cord tissue sections at 35 dpi were permeabilized and blocked as described in “Neutrophil counting”. The sections were then labeled with rat anti-F4/80 (clone BM8, 1:100 dilution, Biolegend) and chicken anti-GFAP (1:200 dilution, Encor). The sections were then washed 3 times for 15 min in 1X PBS and stained with AlexaFluor 555 (1:500, Invitrogen) and AlexaFluor 488 (1:500, JacksonImmuno) conjugated secondary antibodies. After staining, the slides were coverslipped and imaged on a Leica DM6B microscope with a 10X objective. A Matlab code was generated to quantify F4/80^+^ pixels for macrophages and GFAP^+^ pixels for reactive gliosis. The section that contained the highest number of F4/80^+^ pixels was determined to be the lesion epicenter.

### Flow Cytometry Analysis

Mice were anesthetized using 2.5% Avertin (20 μl/g). The peripheral blood was obtained through cardiac puncture using an EDTA (3.6 mg/ml) primed syringe to prevent clotting and placed on ice. Approximately 5 mm of the spinal cord centered on the injury site was dissected and placed in 1X HBSS on ice. All samples were on ice for the entire duration of the flow cytometry staining protocol. The spinal cord tissue was manually dissociated with a plastic tissue pestle and filtered through a 100 μm mesh cell strainer. Red blood cell (RBC) lysis was performed on the blood and spinal cord samples for 5-6 minutes on ice. After RBC lysis, the samples were quenched using NDS-FACS buffer (2% donkey serum and 0.5 μM EDTA in HBSS). After centrifuging and aspirating the supernatant, the pellet was washed with HBSS and stained with Zombie NIR Live/Dead Dye (1:500 dilution, Biolegend) for 30 minutes in the dark. During and after Live/Dead staining, all samples were protected from light. The samples were then washed with NDS-FACS buffer and transferred to a 96 well round bottom plate. Fc blocking was performed using anti-mouse CD16/32 antibody (1:100 dilution, Clone 93, Biolegend) for 20 minutes. Samples were washed in NDS-FACS buffer, partitioned for isotype, surface, and intracellular staining. After separating, the cells were stained in a cocktail of cell surface markers in NDS-FACS buffer for 30 minutes. The following antibodies were used for cell surface staining: Pacific Blue-Ly6G (Clone 1A8, 1:200 dilution, Biolegend), FITC-CD62L (Clone Mel-14, 1:50 dilution, Biolegend), PerCp/Cy5.5-CD11b (Clone M1/70, 1:200 dilution, Biolegend), PE/Cy7-CD101 (Moushi101, 1:100 dilution, Invitrogen) and APC-CD45 (Clone 30-F11, 1:200 dilution, Biolegend). For CD62L isotype staining, the following antibodies were used: Pacific Blue-Ly6G (Clone 1A8, 1:200 dilution, Biolegend), FITC-Rat IgG2a, κ Isotype control (RTK2758, 1:50 dilution, Biolegend), PerCp/Cy5.5-CD11b (Clone M1/70, 1:200 dilution, Biolegend), PE/Cy7-CD101 (Moushi101, 1:100 dilution, Invitrogen) and APC-CD45 (Clone 30-F11, 1:200 dilution, Biolegend). The following antibodies were used for samples for intracellular staining: Pacific Blue-Ly6G (Clone 1A8, 1:200 dilution, Biolegend), PerCp/Cy5.5-CD11b (Clone M1/70, 1:200 dilution, Biolegend), PE/Cy7-CD101 (Moushi101, 1:100 dilution, Invitrogen) and APC-CD45 (Clone 30-F11, 1:200 dilution, Biolegend). After antibody staining, the samples were washed with NDS-FACS buffer and the samples that were not intended for intracellular staining were resuspended in NDS-FACS for same day flow cytometry analysis.

For intracellular staining, samples were washed in NDS-FACS buffer and fixed using CytoFast Fix/Perm Buffer set (Biolegend) for 20 minutes according to the manufacturer’s protocol. After fixation, the samples were washed in permeabilization wash buffer and partitioned into four equal parts. The samples were then incubated in primary antibodies for intracellular staining for 30 minutes. The samples were incubated in one of the following primary antibodies: goat anti-MRP8 (1:2000 dilution, R&D Systems), rabbit anti-CitH3 (1:2000, Abcam) or no primary for staining controls. Samples were washed twice in NDS-FACS and then incubated with the appropriate Alexa-Flour 488-conjugated secondary antibody (anti-rabbit, 1:2000, Invitrogen; anti-goat, 1:2000 Jackson ImmunoResearch) for 30 minutes. Samples were then washed twice with NDS-FACS buffer and resuspended for flow cytometry analysis.

For spinal cord samples, 10 μl of AccuCheck Beads (Invitrogen) was added prior to analysis. Compensation beads were created using Abc Anti-Rat/Hamster Compensation Beads (Invitrogen) for all antibodies and ArC Amine Reactive Compensation beads (Invitrogen) for the Zombie NIR Live/Dead Dye. Flow cytometry was performed using a BD Fortessa X-20. Flow cytometry analysis was performed using the FlowJo Software.

### ELISA

Following collection and centrifugation of the blood and dissociated spinal cord as described in “Flow cytometry analysis”, the supernatant was collected and stored at - 20°C. To determine L-selectin shedding, a soluble L-selectin/CD62L ELISA (DuoSet, R&D Systems) was performed. To determine the extent of neutrophil degranulation, an MRP8/s100a8 ELISA (DuoSet, R&D Systems) was performed. Briefly, a total of 100 μl of supernatant with diluent buffer was incubated overnight at 4°C on NunC plates pre-coated with the appropriate capture antibody. After incubation, the plates were washed with washing buffer and then incubated in 100 μl of the appropriate biotinylated detection antibody for 2 hours at RT according to the manufacturer’s protocol. The plates were then incubated in 100 μl of Streptavidin-HRP for 20 minutes at RT in the dark. Then the plates were incubated in 100 μl of 1X TMB solution (Invitrogen) for 20 – 30 minutes. The reaction was then stopped, and the absorbance of each well was determined using a microplate reader (Molecular Devices, SpectraMax) set to 450 nm.

To determine neutrophil extracellular trap formation, an enzyme-DNA capture ELISA was performed. A total of 50 μl of supernatant with diluent buffer was incubated overnight at 4°C on NunC plates pre-coated with anti-CitH3 antibody (1:250, Abcam). After incubation, the plates were washed and incubated in 50μl of anti-DNA-POD (1:100, clone MCA-333, Roche) for 2 hours at RT. Then the plates were incubated in 100 μl of 1X TMB solution (Invitrogen) for 20 minutes. The reaction was then stopped, and the absorbance of each well was determined using a microplate reader (Molecular Devices, SpectraMax) set to 450 nm.

### Western blotting

Protein for Western blotting was isolated from the spinal cord at 1 dpi. Mice were anesthetized using 2.5% Avertin and 5 mm of the spinal cord centered at the lesion epicenter was dissected and placed in lysis buffer (RIPA buffer with protease inhibitor). The samples were on ice for the duration of the sample preparation protocol. The spinal cord tissue was manually dissociated using a plastic tissue pestle. Homogenates were then incubated in the lysis buffer on an orbital shaker for 60 minutes. After 60 minutes, the samples were centrifuged at 12000 rpm at 4°C for 15 minutes. The supernatant was collected, aliquoted, and the protein concentration of each sample was determined using the Bicinchoninic Acid Assay (Pierce BCA Protein Assay Kit, Thermo Fischer). A total of 156 μg of the sample was added to LDS nuPAGE reaction master mix (4X LDS, 2x Beta-mercaptoethanol, DI water) at a 1:1 ratio. The samples were boiled for 10 minutes at 95°C and stored at −20°C.

Western blots were performed on pre-made 10% Bis-Tris gels (ThermoFisher). A total of 20 μg of sample protein was loaded into each well and 2 μl of the ladder (ThermoFisher, LC5925) was added on the first and last well of the gel. The samples were run in NuPAGE MES SDS Running Buffer (ThermoFisher) for 90 minutes at 125V. The gel was placed in the transfer stack according to the manufacturer’s guidelines (iBlot 2 Dry Blotting System, Invitrogen) and was transferred to the membrane for 6 minutes at 25V. Following the transfer, the membrane was cut down to size and blocked for 1 hour at room temperature in tris-buffered saline (TBS) + 5% bovine serum albumin (BSA) to prevent non-specific binding.

Hemoglobin (Rabbit anti-mouse,1:5000, ThermoFisher) primary and IgG secondary antibodies (Donkey anti-mouse, IRDye 800CW, 1:15000, LI-COR) were diluted in TBST (TBS + 0.1% Tween20) + 5% BSA and incubated with the membrane blot overnight at 4°C on a rocker. The following day, the blots were washed 3 times using TBST for 10 minutes. The secondary antibody (Donkey anti-Rabbit IgG, IRDye 680RD, 1:1500, LI-COR) was diluted in TBST + 5% BSA and incubated with the blots for 1 hour at RT on a rocker. After secondary antibody incubation, the blots were washed 3 times using TBST for 10 minutes. Before imaging, the blots were washed and transported in TBS. The blots were imaged on an Odyssey CLx Imager (Li-Cor). After imaging, the blots were then incubated in GAPDH (Rabbit anti-mouse, 1:5000, Millipore) primary antibody overnight at 4°C on a rocker followed by secondary antibody (Donkey anti-Rabbit IgG, 1:1500, LI-COR) incubation. The blots were then imaged again on a Li-Cor Odyssey CLx Imager.

### Experimental Design and Statistical Analysis

Throughout the behavioral studies, the surgeon and observers were blinded to the animal’s genotype to minimize bias in the surgical procedure and data collection. Each animal was assigned a unique experimental identification number to ensure the observers remained unaware of the genotype or the experimental condition. Simple or block randomization was employed for the studies.

The statistical analysis was performed using GraphPad Prism. A two-way or three-way repeated measures ANOVA followed by a Sidak’s multiple comparisons test was used to evaluate the BMS scores. White matter sparing and neutrophil accumulation spanning the injury site were evaluated with a three-way ANOVA followed by a Tukey’s post hoc test. If values were missing, the mixed effects model was used in place of repeated measures. White matter sparing and neutrophil accumulation at the lesion epicenter were evaluated with an unpaired two-tailed Student’s *t* test or a two-way ANOVA followed by a Tukey’s post hoc test. ELISA and western blot were evaluated with a two-way ANOVA followed by a Tukey’s post hoc test. Flow cytometry where two tissues from the same animal were assessed were evaluated with a three-way ANOVA. Neutrophil clearance across time points was evaluated using a three-way ANOVA mixed-effects analysis. At 35dpi, neutrophil clearance was assessed using a two-way ANOVA followed by Tukey’s post hoc test. Simple linear regression analysis was performed to assess the relationship between neutrophil accumulation and BMS scores. Statistical significance is defined as p<0.05. Data are represented as mean ± SEM.

## Results

### L-selectin mediates long-term recovery and tissue sparing in a sex dependent manner

To investigate the role of endogenous L-selectin shedding in functional recovery after SCI, we compared the hindlimb locomotor recovery using the Basso Mouse Scale (BMS) in WT and L(E) male and female mice following a moderate thoracic contusion SCI. Interestingly, we observed a sex-dependent effect of L-selectin shedding on long-term recovery in the BMS main and sub-scores (Figure 1A-B). When data were disaggregated by sex and further analyzed, we found that L(E) female mice had reduced hindlimb functional recovery when compared to WT females. However, there was no difference in long-term functional recovery between WT and L(E) male mice (Figure 1C-D).

**Figure 1.**
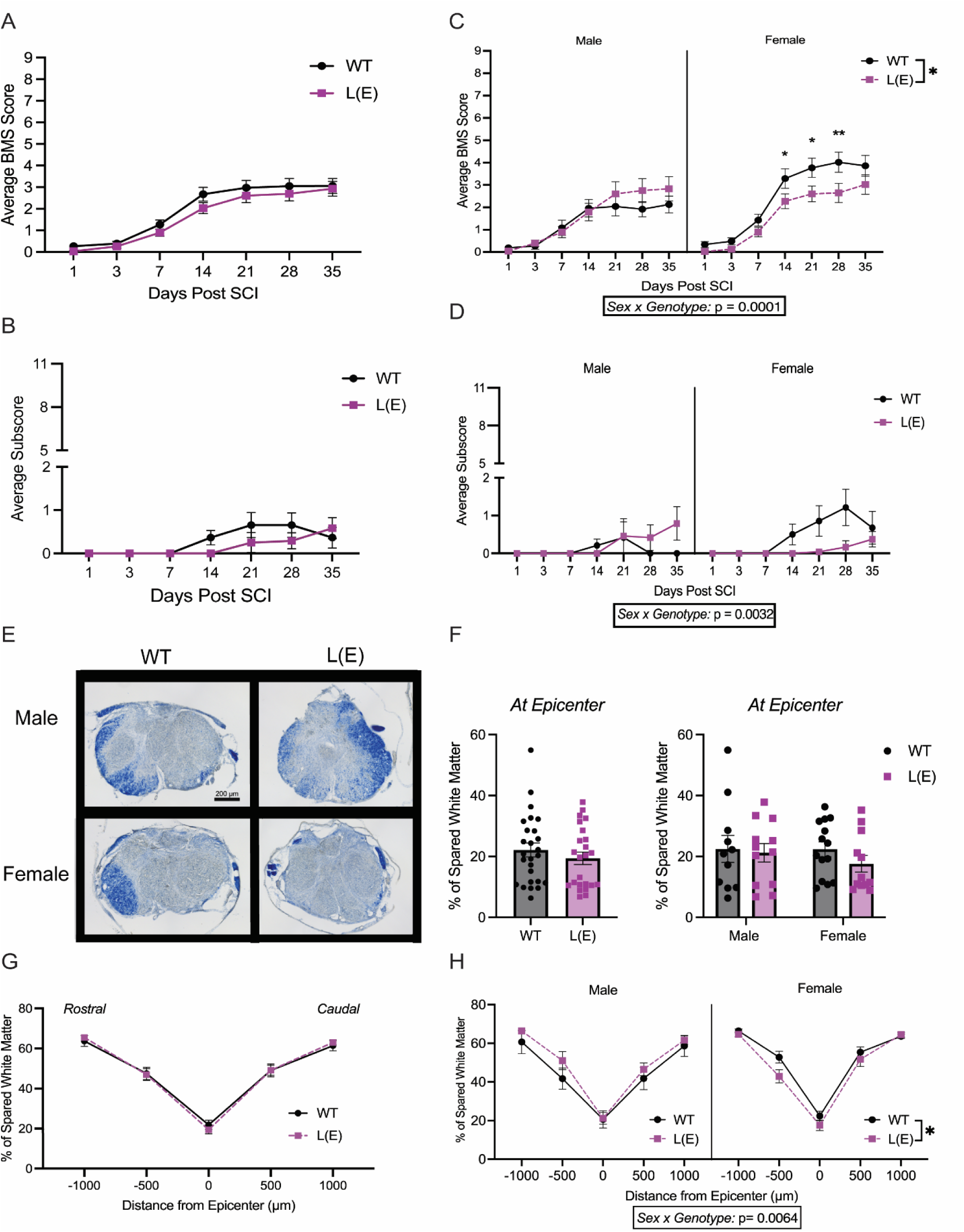
Genetic loss of L-selectin shedding impairs functional recovery and tissue sparing in a sex-dependent manner. **(A-D)** Long-term hindlimb recovery after SCI was assessed using the BMS. Loss of L-selectin shedding alters long-term recovery in a sex-dependent manner. Disaggregation and analysis of data by sex demonstrates that long-term recovery is impaired in L(E) female mice when compared to WTs, but not males. (**E)** Representative images of white matter sparing at the lesion epicenter in WT and L(E) male and female mice. **(F)** There is no difference in the percentage of white matter spared at the lesion epicenter between WT and L(E) mice in either sex. **(G&H)** Loss of L-selectin shedding alters the percentage of white matter spared throughout the lesion in a sex-dependent manner. Disaggregation and analysis of data by sex demonstrates that white matter sparing is reduced in L(E) female mice relative to WTs. 3-way repeated measures ANOVA followed by 2-way repeated measures ANOVA with Sidak’s post-hoc test for A, B, C, D, G and H; Student’s t-test and 2-way ANOVA for F. n = 11-14/sex/genotype, data are shown as mean ± SEM. * p<0.05, ** p<0.01, ***p<0.001, p<****0.0001. Abbreviations: BMS = Basso Mouse Scale, dpi = days post-injury

We next investigated the role of L-selectin shedding in white matter tissue sparing following SCI. We observed no difference in the percentage of white matter spared at the injury epicenter between WT and L(E) mice in either sex (Figure 1F). However, when we examined the spared white matter for 2 mm across the SCI lesion, we observed a sex-dependent effect of L-selectin shedding on total white matter spared (Figure 1G-H). When the data were disaggregated by sex and further analyzed, we observed reduced white matter sparing throughout the injury site only in L(E) female mice relative to WTs, but not males. Collectively, these data show that genetic loss of endogenous L-selectin shedding impairs long-term recovery and tissue sparing in a sex-dependent manner.

### L-selectin shedding regulates acute neutrophil accumulation in a sex-dependent manner

L-selectin is known to be involved in the tethering and rolling of neutrophils on the inflamed endothelium during migration towards the site of injury or infection^30,31^. To investigate if L-selectin shedding alters neutrophil accumulation after SCI, we performed immunostaining and semi-automated counting of Ly6G^+^ neutrophils in transverse tissue sections spanning the injury site (Figure 2A). We observed no differences in the number of neutrophils at the injury epicenter between WT and L(E) mice in either sex (Figure 2B). However, when we analyzed the number of neutrophils across the lesion, we found that endogenous L-selectin shedding alters neutrophils accumulation at 1 dpi in a sex-dependent manner. Specifically, we observed a trend towards lower neutrophil numbers in L(E) female mice when compared to WT females and the opposite pattern was observed in male mice (Figure 2C). Similar sex and genotype-dependent effects were observed for neutrophil accumulation in the grey and white matter of the injured spinal cord, but not the meninges (Figure S1). These data indicate that L-selectin shedding affects neutrophil accumulation in a sex-dependent manner.

**Figure 2.**
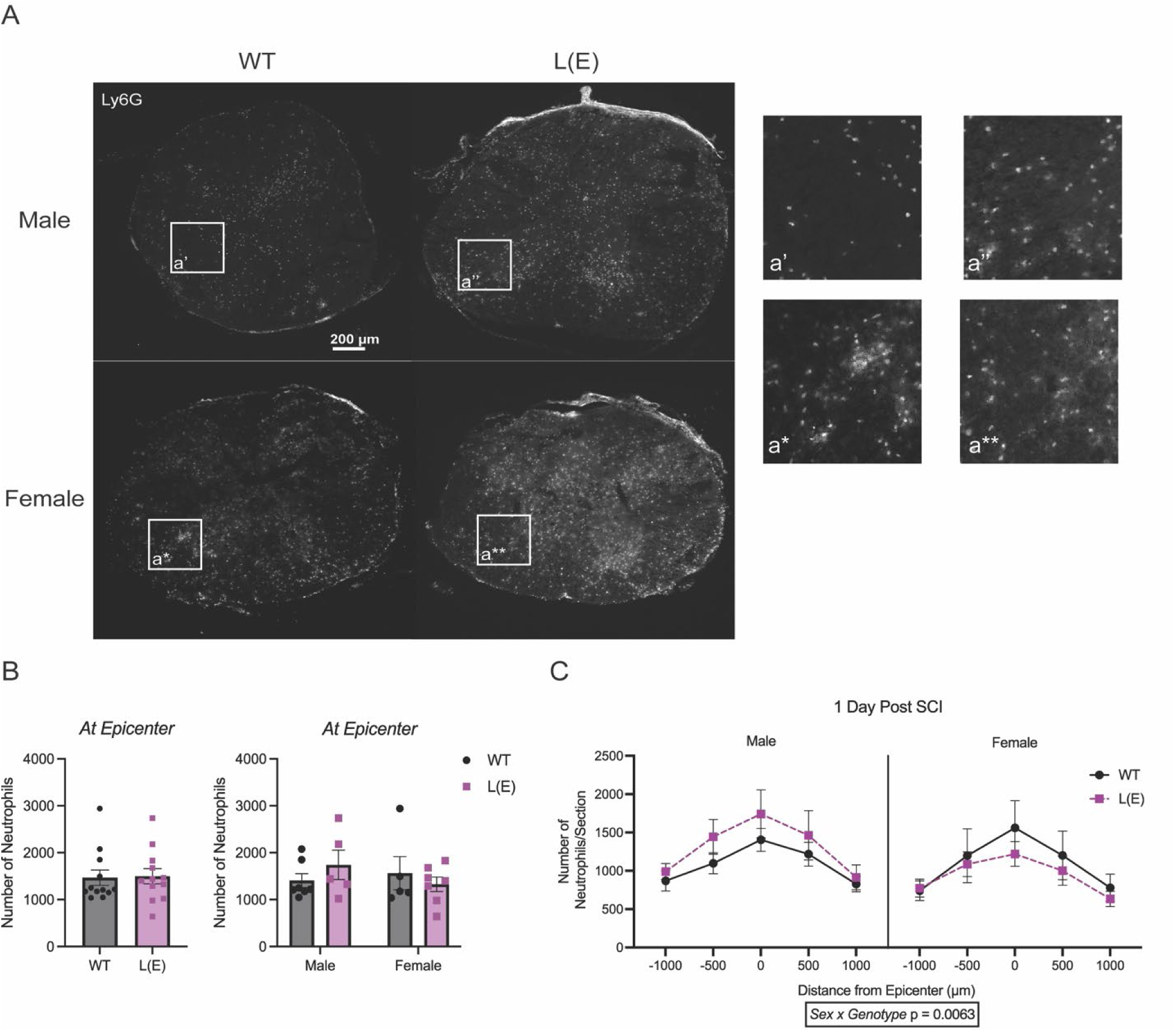
L-selectin shedding regulates neutrophil accumulation in a sex-dependent manner. **(A)** Representative images of Ly6G^+^ neutrophils at the lesion epicenter of WT and L(E) male and female mice. **(B)** Neutrophil accumulation at the injury epicenter at 1 dpi. No differences were observed between L(E) and WT mice in either sex. **(C)** L-selectin shedding alters total neutrophil accumulation across the SCI lesion in a sex-dependent manner. Students t-test and 2-way ANOVA for A, 3-way repeated measures ANOVA for B, n = 5-7/sex/genotype, data are shown as mean ± SEM. * p<0.05, ** p<0.01, ***p<0.001, p<****0.0001. Abbreviations: dpi = days post-injury

### L-selectin shedding does not mediate neutrophil activation or effectors functions acutely after SCI

Since we observed that L-selectin shedding influences neutrophil accumulation in a sex-dependent manner, we next aimed to investigate other neutrophil responses in the acutely injured spinal cord. We first examined L-selectin shedding by analyzing soluble L-selectin in the supernatant of blood and spinal cord samples, as well as activation markers (L-selectin and CD11b) on the surface of circulating and intraspinal neutrophils at 1 dpi. Soluble L-selectin is commonly found in the blood of both mice and humans^50–52^, potentially due to gradual L-selectin shedding during neutrophil aging^53^. Interestingly, we observed that WT mice had higher levels of soluble L-selectin in the blood compared to L(E) mice, demonstrating impaired L-selecting shedding. As anticipated, we also found higher levels of soluble L-selectin in the spinal cord of WT mice compared to L(E) mice (Figure 3A&B). To verify L-selectin shedding on neutrophils, we examined CD62L G-MFI (geometric mean fluorescence intensity) on the surface of neutrophils by flow cytometry. While we observed no differences in L-selectin levels on circulating neutrophils in WT or L(E) mice, intraspinal neutrophils from WT mice had markedly reduced L-selectin compared to L(E) mice as expected (Figure 3C&D). To determine if soluble L-selectin could be coming from other leukocytes in high abundance at 1 dpi, we examined L-selectin expression on microglia. We observed no differences in CD62L G-MFI on microglia from uninjured WT and L-selectin KO mice, indicating that L-selectin is not normally expressed on microglia (Figure S2). Since neutrophil activation cannot be assessed by L-selectin shedding in L(E) mice, we analyzed the levels of CD11b (integrin alpha-M), which increases on the cell surface upon activation^23^. While we observed an increase in CD11b levels on intraspinal neutrophils relative to their circulating counterparts, there were no difference in the levels of CD11b between L(E) and WT mice in either sex (Figure 3 E&F). Collectively, these data demonstrate that while L-selectin shedding is impaired on neutrophils from L(E) mice after SCI, neutrophil activation does not appear to be affected.

**Figure 3.**
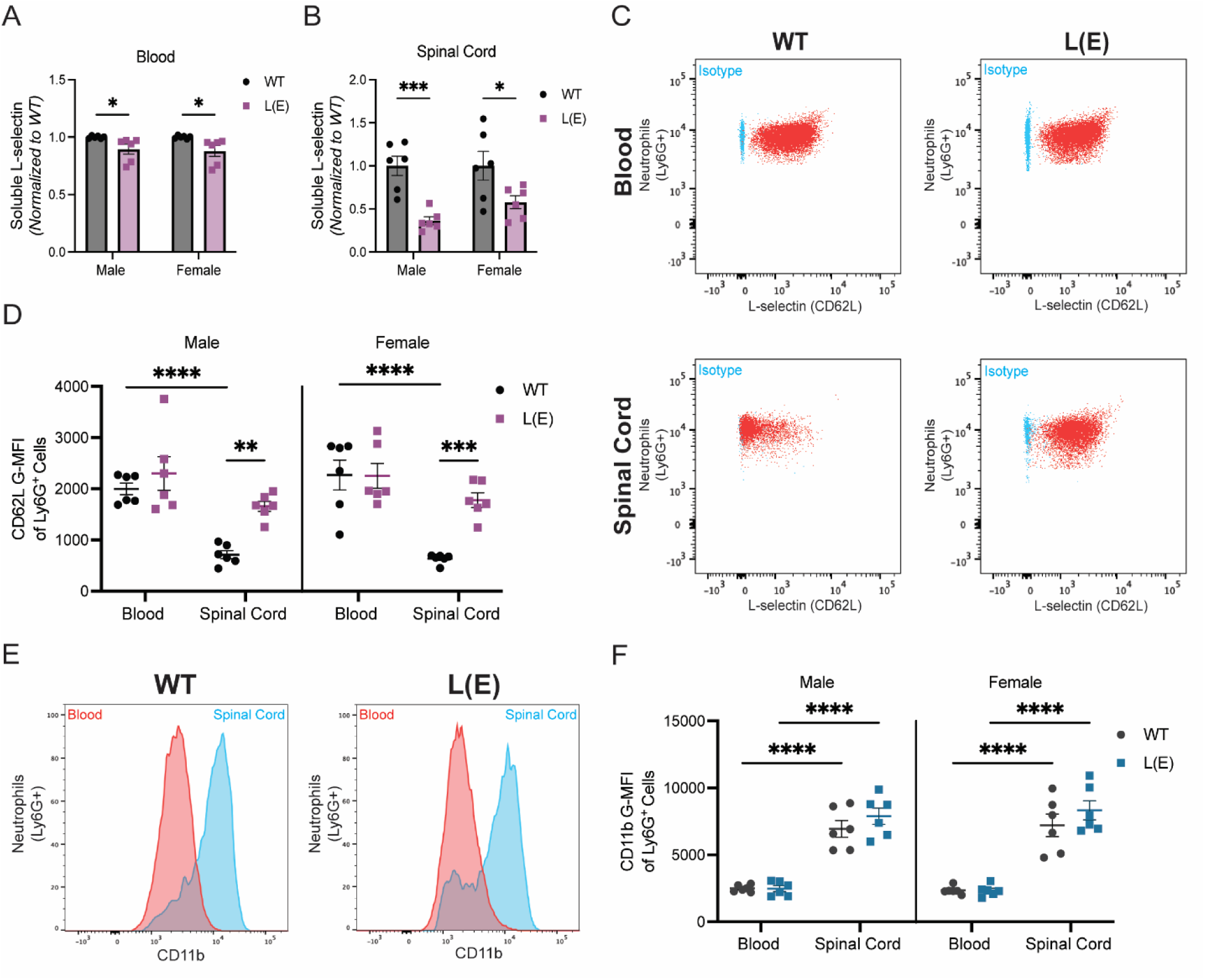
L-selectin shedding does not mediate neutrophil activation in the injured spinal cord. **(A&B)** Soluble L-selectin is reduced in L(E) mice in both the blood and spinal cord at 1 dpi. **(C)** Representative flow cytometry dot plots of L-selectin (CD62L) levels on WT and L(E) neutrophils from the blood and spinal cord. **(D)** Quantification of G-MFI L-selectin levels are the same on neutrophils in the blood of WT and L(E) mice but reduced only on intraspinal neutrophils from WT mice. **(E)** Representative flow cytometry histograms of CD11b levels on WT and L(E) neutrophils from the blood and spinal cord. **(F)** CD11b G-FMI is similar in the blood and spinal cord of L(E) mice relative to WTs, indicating no difference in neutrophil activation. 2-way ANOVA for A and B, 3-way ANOVA for D and F, n = 6/sex/genotype, data are shown as mean ± SEM. * p<0.05, ** p<0.01, ***p<0.001, p<****0.0001. Abbreviations: dpi = days post-injury, G-MFI = geometric mean fluorescent intensity

We next analyzed neutrophil effector functions, including degranulation and NET formation, in the blood and acutely injured spinal cord. We observed no difference in the amount of MRP8, a granular protein released during degranulation, in the supernatant of the blood and spinal cord at 1 dpi between L(E) or WT mice (Figure 4A). While we did observe an increase in degranulation of intraspinal neutrophils relative to their circulating counterparts (Figure 4B), there were no differences in degranulation of neutrophils between L(E) and WT mice. Additionally, we observed no differences in the amount of NET fragments, measured by CitH3 and DNA complexes, in the supernatant from the blood and spinal cord from L(E) mice relative to WTs (Figure 4C). Similar to degranulation, we did observe an increase in the percentage of intraspinal neutrophils undergoing NET formation relative to circulating neutrophils (Figure D). However, there were no differences in CitH3^+^ neutrophils between L(E) and WT mice. To delve deeper into the characteristics of infiltrating neutrophils, we analyzed their maturation profiles using CD101. We observed no differences in neutrophil maturity in the blood or spinal cord between L(E) and WT mice (Figure S4). Altogether, our data demonstrate that neutrophil activation, maturation, and effector functions in the acutely injured spinal cord do not appear to be affected by L-selectin shedding.

**Figure 4.**
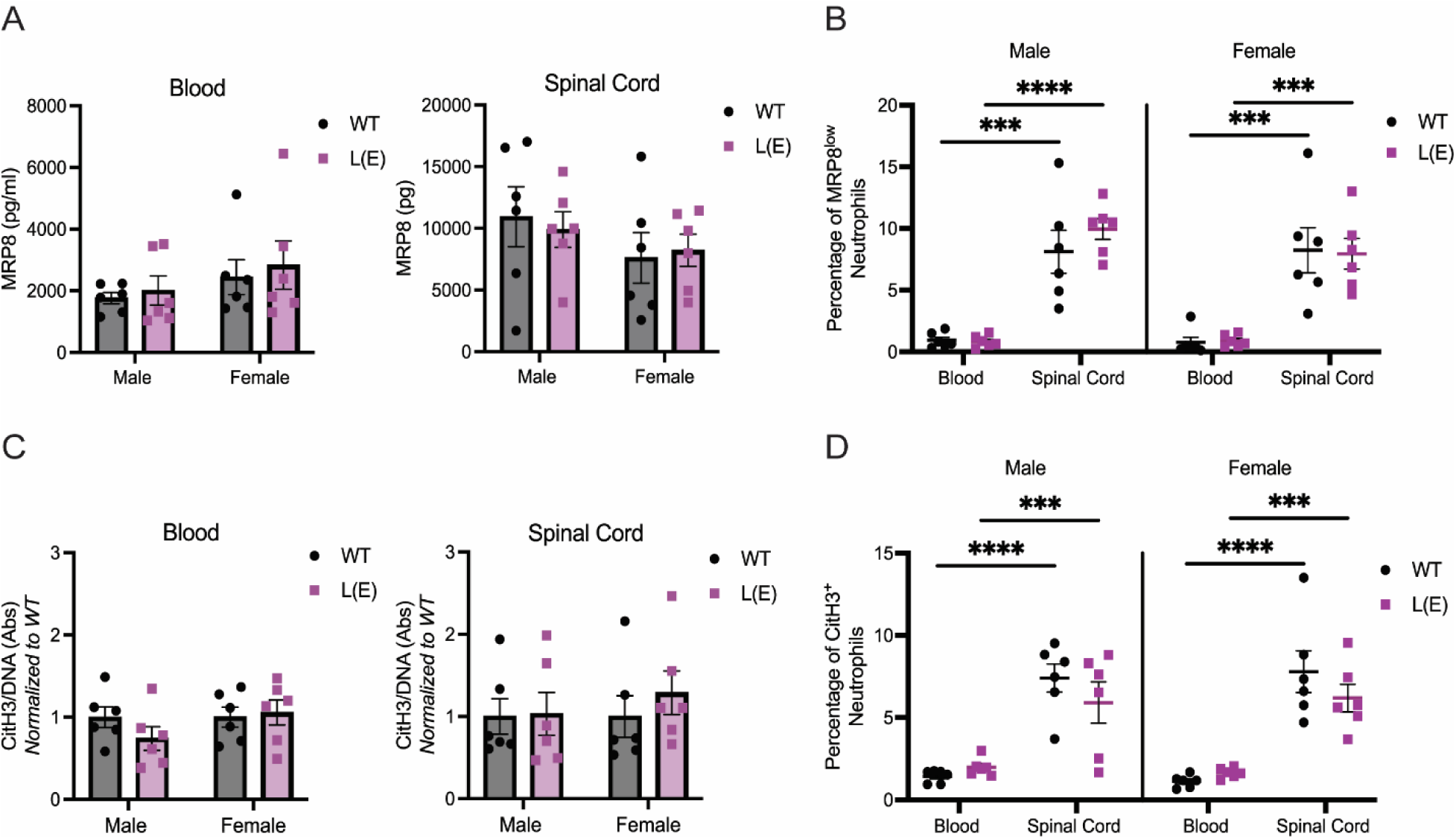
L-selectin shedding does not mediate neutrophil effector functions in the injured spinal cord. **(A)** MRP8 levels are similar in the supernatant from the blood and spinal cord of WT and L(E) mice at 1dpi. **(B)** The percentage of neutrophils undergoing degranulation (MRP8^low^) in the spinal cord is increased relative to the blood. No difference was observed between L(E) and WT mice. **(C)** There is no difference in CitH3 and DNA complexes found in the supernatant blood and spinal cord of WT and L(E) mice. **(D)** Neutrophil extracellular trap formation increases in intraspinal neutrophils relative to the blood at similar levels in WT and L(E) mice. 2-way ANOVA for A and C, 3-way ANOVA for B and D, n = 6/sex/genotype, data are shown as mean ± SEM. ***p<0.001, p<****0.0001. Abbreviations: dpi = days post-injury

### L-selectin shedding does not affect acute blood spinal cord barrier disruption

We explored the role of L-selectin shedding in the disruption of the blood-spinal cord barrier (BCSB), as neutrophils have been implicated in mediating BSCB damage during neuroinflammatory diseases^54,55^. We characterized both hemorrhage and BCSB permeability at 1 dpi in spinal cord tissue homogenates by Western blot analysis for hemoglobin (as a surrogate for red blood cells) and extravasated endogenous IgG (Figure 5A). We observed no differences in hemorrhage in the acutely injured spinal cord between L(E) and WT mice (Figure 5B). Additionally, we found no difference in the amount of IgG heavy and light chain between L(E) and WT mice indicating that endogenous L-selectin shedding does not alter disruption of the BSCB (Figure 5C&D). However, we did notice that female mice had significantly higher IgG levels compared to males, suggesting potential sex differences in secondary injury after SCI (IgG *heavy chain* p_sex_=0.0151, IgG *light chain* p_sex_=0.230).

**Figure 5.**
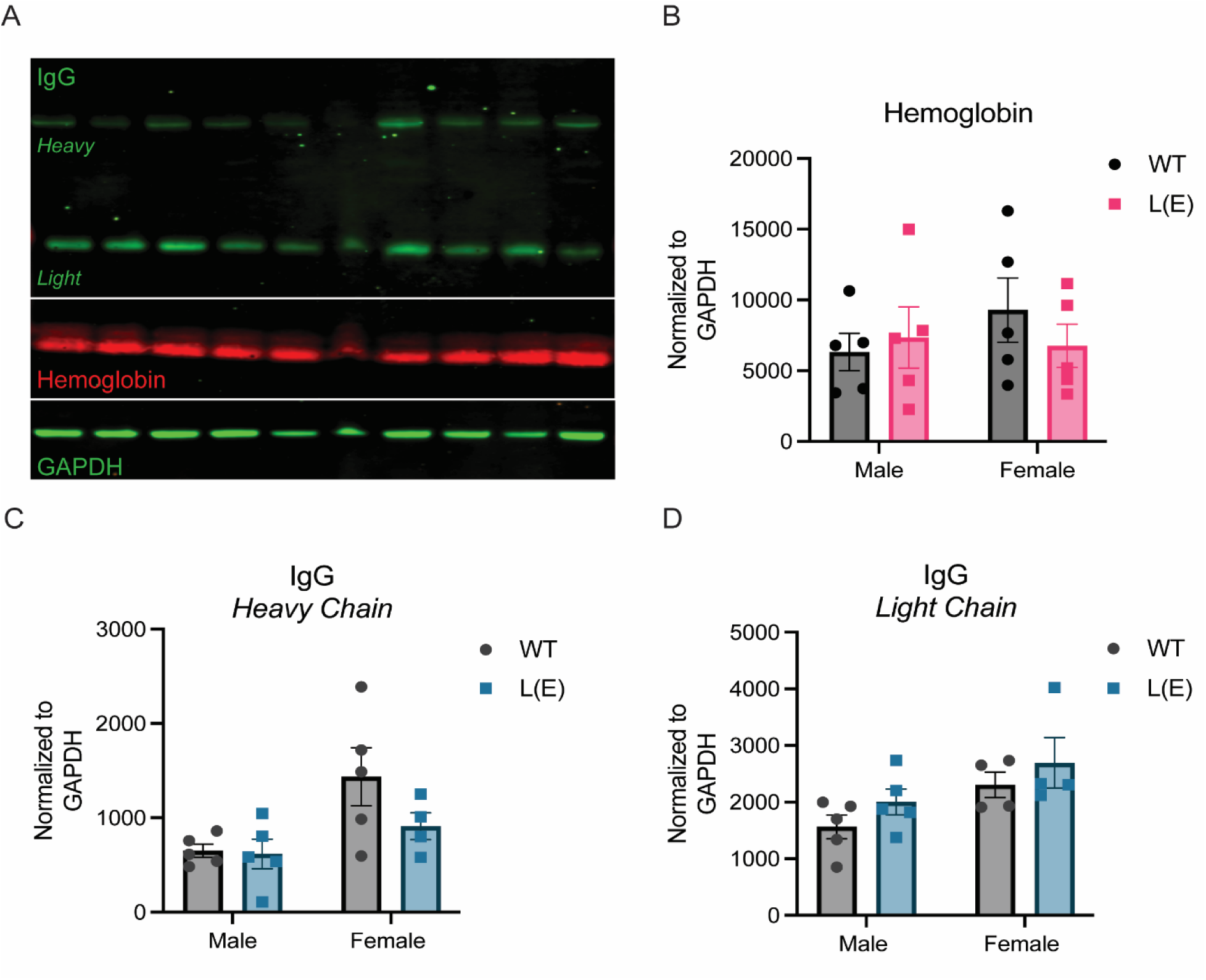
L-selectin shedding does not alter acute hemorrhage or permeability of the BSCB after SCI. **(A)** Representative image of the Western blot for IgG, Hemoglobin, and GAPDH (internal control). **(B)** At 1 dpi, there was no difference in the amount of accumulated hemoglobin, indicating hemorrhage, in the spinal cord in WT and L(E) mice. **(C & D)** There is no difference in the amount of accumulated IgG, indicating BSCB permeability, in the spinal cord between WT and L(E) mice. Higher levels of IgG were found in female mice relative to males (IgG *heavy chain* p_sex_=0.0151, IgG *light chain* p_sex_=0.230). 2-way ANOVA, n = 4-5/sex/genotype, data are shown as mean ± SEM. Abbreviations: dpi = days post-injury

### L-selectin shedding mediates neutrophil clearance in a sex-dependent manner

Given that L-selectin shedding mediates sex-dependent differences in long-term functional recovery but does not appear to alter acute neutrophil function and secondary injury, we next investigated long-term neutrophil clearance after SCI. Neutrophils typically peak by 1 dpi in rodent models of SCI and are thought to be cleared by efferocytosis or rTEM. Thus, we examined neutrophil numbers in the injured spinal cord at 3, 14, and 35 dpi. Similar to 1 dpi, we observed a sex- and genotype-dependent difference in neutrophils distributed across the injury site at 3 dpi (Figure 6A & S5). At 14 and 35 dpi, however, no differences were observed between L(E) and WT mice in either sex (Figure 6B-C). Considering the differences in initial neutrophil accumulation at 1 dpi, we normalized neutrophil numbers to 1 dpi and found that L(E) female mice trended towards greater retention of neutrophils across time compared to WT females (Figure 7A). No differences were observed in males. At 35 dpi, L(E) female mice had a higher relative number of neutrophils persisting compared to WT females (Figure 7B). In L(E) female mice, but not WT females or male mice, we found that the number of neutrophils at the injury epicenter significantly correlated with the average BMS score at 35 dpi (Figure 7C). To determine if L-selectin shedding affects other long-term injury inflammatory processes, we assessed total macrophage accumulation and reactive gliosis at 35dpi. We did not observe any difference in the accumulation of F4/80^+^ macrophages or GFAP levels in the injured spinal cords of WT and L(E) mice (Figure S6). Take together, these data suggest that L-selectin shedding mediates neutrophil clearance in a sex-dependent manner, which corresponds with long-term functional recovery.

**Figure 6.**
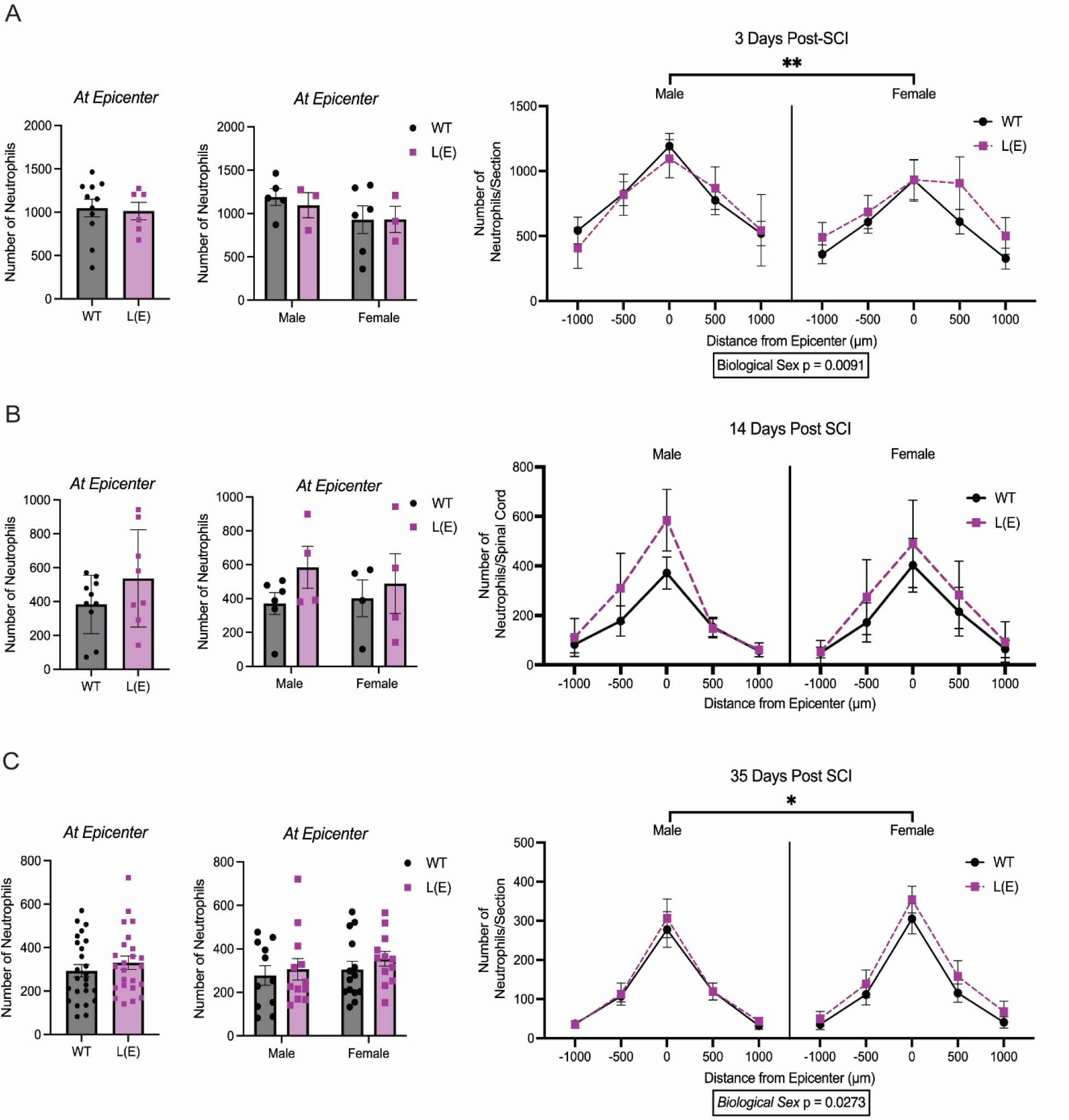
L-selectin shedding alters longer term neutrophil accumulation in a sex-dependent manner. **(A)**. No differences were observed in neutrophil accumulation at the injury epicenter at 3 dpi between L(E) and WT mice in either sex. However, L-selectin shedding alters total neutrophil accumulation across the SCI lesion at 3 dpi in a sex-dependent manner (n = 3 – 6/sex/genotype). **(B)** L-selectin shedding does not alter the accumulation of neutrophils at 14 dpi (n = 4 – 7/sex/genotype). **(C)** No differences were observed in neutrophil accumulation at the injury epicenter at 35 dpi between L(E) and WT mice in either sex. However, L-selectin shedding alters total neutrophil accumulation across the SCI lesion at 35 dpi in a sex-dependent manner (n = 11 – 14/sex/genotype) Students t-test, 2-way ANOVA, 3-way ANOVA, data are shown as mean ± SEM. * p<0.05, ** p<0.01. Abbreviations: dpi = days post-injury

**Figure 7.**
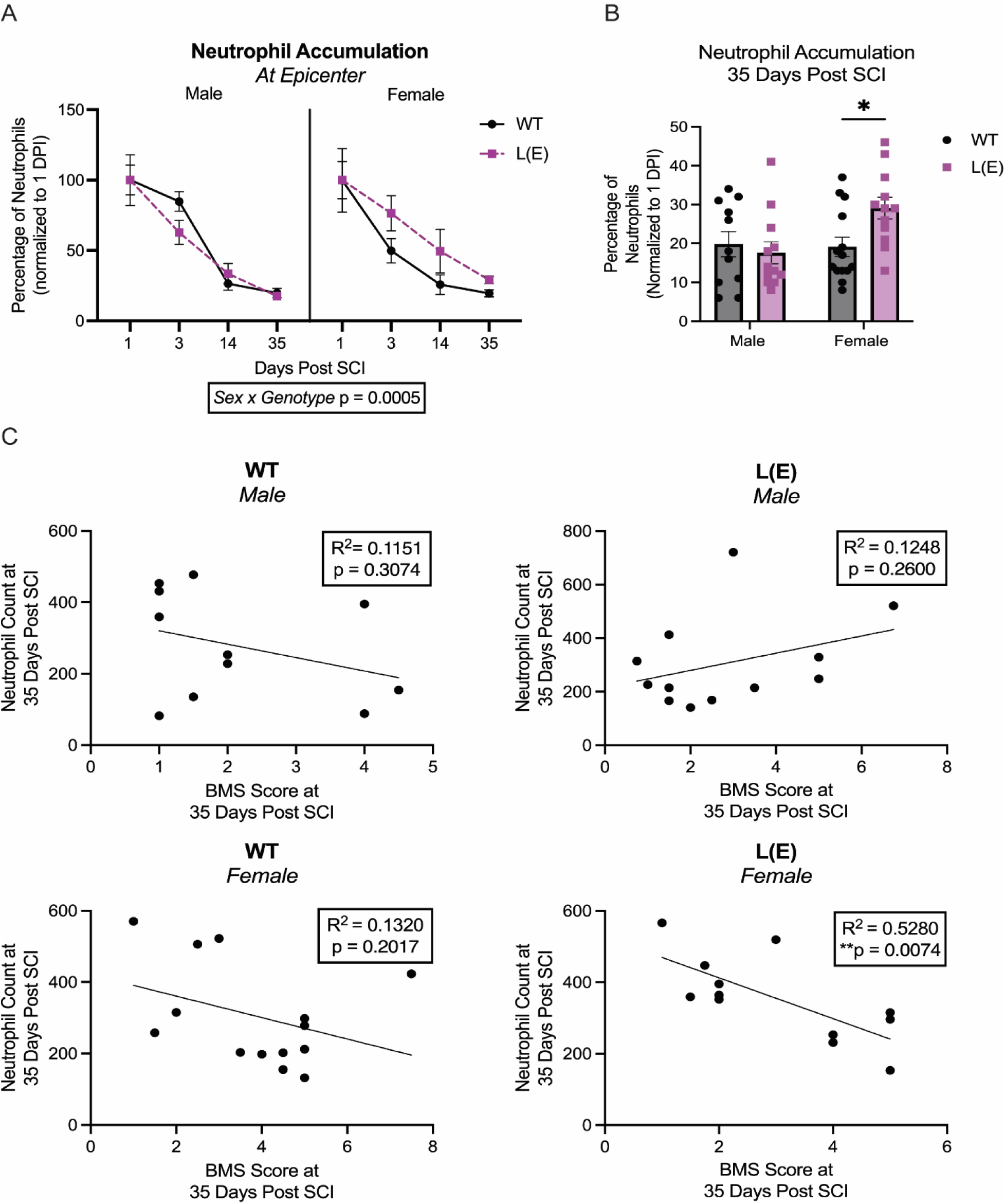
L-selectin shedding regulates neutrophil clearance in a sex-dependent manner. **(A&B)** Neutrophil abundance in each group was normalized to 1 dpi. L-selectin shedding mediates neutrophil clearance in a sex-dependent manner. Specifically, L(E) female mice retained a higher percentage of neutrophils at the injury epicenter when compared to WT. No differences were observed in males. **(C)** The number of neutrophils at the injury epicenter correlated with the average BMS score in L(E) female mice, but not WT females or males. 2-way ANOVA, 3-way ANOVA, Linear regression analysis, n = 11 – 14/sex/genotype, data are shown as mean ± SEM. * p<0.05, ** p<0.01.

## Discussion

We have previously shown that L-selectin is a critical mediator of secondary injury and long-term neurologic deficits after SCI. We also found that augmenting L-selectin shedding early after SCI improves recovery and tissue sparing in male mice^47^. However, the role of endogenous L-selectin shedding in long-term recovery and neutrophils responses after SCI has yet to be investigated, particularly when also considering sex as a biological variable. Here, we examined how genetic abrogation of endogenous L-selectin shedding affects neutrophil accumulation and functions, secondary injury, and long-term outcomes after SCI.

To examine the role of L-selectin shedding in SCI, we utilized L(E) mice in which the non-cleavable membrane proximal domain of E-selectin was knocked into the cleavage site of the L-selectin to prevent shedding by cell surface enzymes such as a disintegrin and metalloproteinases (ADAMs). Utilizing L(E) mice and their WT littermates, we investigated the sex-dependent role of endogenous L-selectin shedding on functional recovery and tissue sparing after SCI. In this study, we found that genetic abrogation of L-selectin shedding impairs long-term recovery in female, but not male mice. Hindlimb functional recovery is typically associated with greater white matter sparing^56^. While the absence of L-selectin shedding does not affect white matter sparing at the lesion epicenter, total white matter spared across the SCI lesion is altered in a sex-dependent manner. Specifically, blocking L-selectin shedding impairs white matter sparing in females alone. Collectively, our findings demonstrate that L-selectin shedding improves long-term outcomes in a sex-dependent manner.

The role of L-selectin in leukocyte responses has most prominently been studied in neutrophils, although many leukocyte subtypes express L-selectin. We have previously shown that genetic deficiency of L-selectin reduces neutrophil accumulation, but not the accumulation of other leukocyte subtypes, into the acutely injured spinal cord^47^. To rule out a potential role for microglia, we examined L-selectin levels on microglia in the uninjured spinal cord. Previous research has found conflicting evidence on whether microglia express L-selectin^57–59^. However, we found that microglia do not express detectable levels of L-selectin prior to SCI and thus likely do not undergo L-selectin shedding upon injury and cellular activation. Neutrophils express high levels of L-selectin in circulation, and we demonstrate that extensive L-selectin shedding occurs on intraspinal neutrophils in WT, but not L(E), mice regardless of sex. Interestingly, we found that L-selectin shedding affects neutrophil accumulation in the SCI lesion in a sex-dependent manner. To the best of our knowledge, our study is the first to demonstrate sex differences in L-selectin shedding-mediated neutrophil accumulation into a site of injury.

L-selectin has been shown to facilitate neutrophil activation and effector functions such as degranulation, NET formation, cytokine expression, and ROS production^28,30,32–39^. However, it is not known whether L-selectin shedding regulates neutrophil activities during CNS injury. We investigated neutrophil activation, degranulation, and NET formation at 1 dpi, which corresponds to the peak of neutrophil accumulation^7,10,11,23,27^. L-selectin is also known to play a role in signaling that leads to the activation of integrins, like CD11b^60^. We found that activation of neutrophils, as assessed by CD11b levels on the cell surface, is not mediated by L-selectin shedding in the context of SCI. Similarly, we also found that L-selectin shedding does not mediate neutrophil degranulation or NET formation as assessed by ELISA and flow cytometry. While neutrophil accumulation is dependent on L-selectin shedding, the subsequent activation of neutrophils and their effector functions in the acutely injured spinal cord appears to be predominantly mediated through other receptors and signaling pathways.

Neutrophils have canonically been considered to mediate inflammatory damage and disruption of the BSCB leading to extravasation of serum proteins and the development of hemorrhage^61,54^. Since we did not observe differences in neutrophil effector functions, we assessed if L-selectin shedding alters acute secondary damage. We found that L-selectin shedding does not alter the extent of hemorrhage or BSCB permeability in either sex. Collectively, our data indicate that L-selectin shedding may mediate later occurring leukocyte responses that impact long-term recovery. We found that L-selectin shedding does not appear to mediate total macrophage abundance or reactive gliosis at 35 dpi, suggesting that L-selectin shedding does not substantially influence monocyte, macrophage, or astrocyte responses to SCI.

As previously reported by other studies, we observed a decrease in neutrophil numbers after 1 day post-SCI^11^. At 3 dpi, we found that neutrophils numbers begin to decrease, but that L-selectin shedding still altered neutrophil numbers in a sex-dependent manner. Neutrophil numbers continued to decline at 14 and 35 dpi, and we observed a trend towards increased neutrophils in the absence of L-selectin shedding in female mice at 35 dpi. Since genetic abrogation of L-selectin shedding appeared to reduce neutrophil accumulation at 1 dpi, we normalized neutrophil counts at the lesion epicenter to 1 dpi. By assessing the relative persistence of neutrophils in each group, we found that L-selectin shedding reduces neutrophil persistence in female mice, but not males, after SCI. L-selectin has been implicated in the clearance of neutrophils from an inflammatory environment^43,45,46^. Our data is the first to demonstrate a sex-dependent role for L-selectin shedding in neutrophil clearance from the injured spinal cord. In L(E) female mice, neutrophil abundance at 35 dpi correlated with BMS scores indicating a potential relationship between neutrophil persistence and long-term recovery. The mechanisms of L-selectin shedding-mediated neutrophil clearance (efferocytosis or rTEM), as well as the neutrophil activities leading to long-term tissue damage, should be investigated in future studies.

In this study, we establish that endogenous shedding of L-selectin is sufficient to improve hindlimb functional recovery in female, but not male, mice. While we show that L-selectin shedding does not alter neutrophil effector functions in the injured spinal cord, we demonstrate that the shedding of L-selectin regulates neutrophil accumulation and clearance in a sex-dependent manner. In summary, our findings further support L-selectin shedding as a promising mechanism to reduce inflammatory damage and improve long-term outcomes after SCI. Future studies should address the therapeutic potential of augmenting L-selectin shedding in female mice and determine the mechanisms driving neutrophil clearance from the injured spinal cord.

## Acknowledgements

The authors would like to thank the Texas A&M University Flow Cytometry & Cell Sorting Facility and the Microscopy and Imaging Center for the use of their equipment. Funding support was provided by Mission Connect, a program of TIRR Foundation, NIH R01NS122961, NSF GRFP 2139772 (MRP), and Wings for Life (SKR).

## Supplemental Figures

**Supplemental Figure 1.**
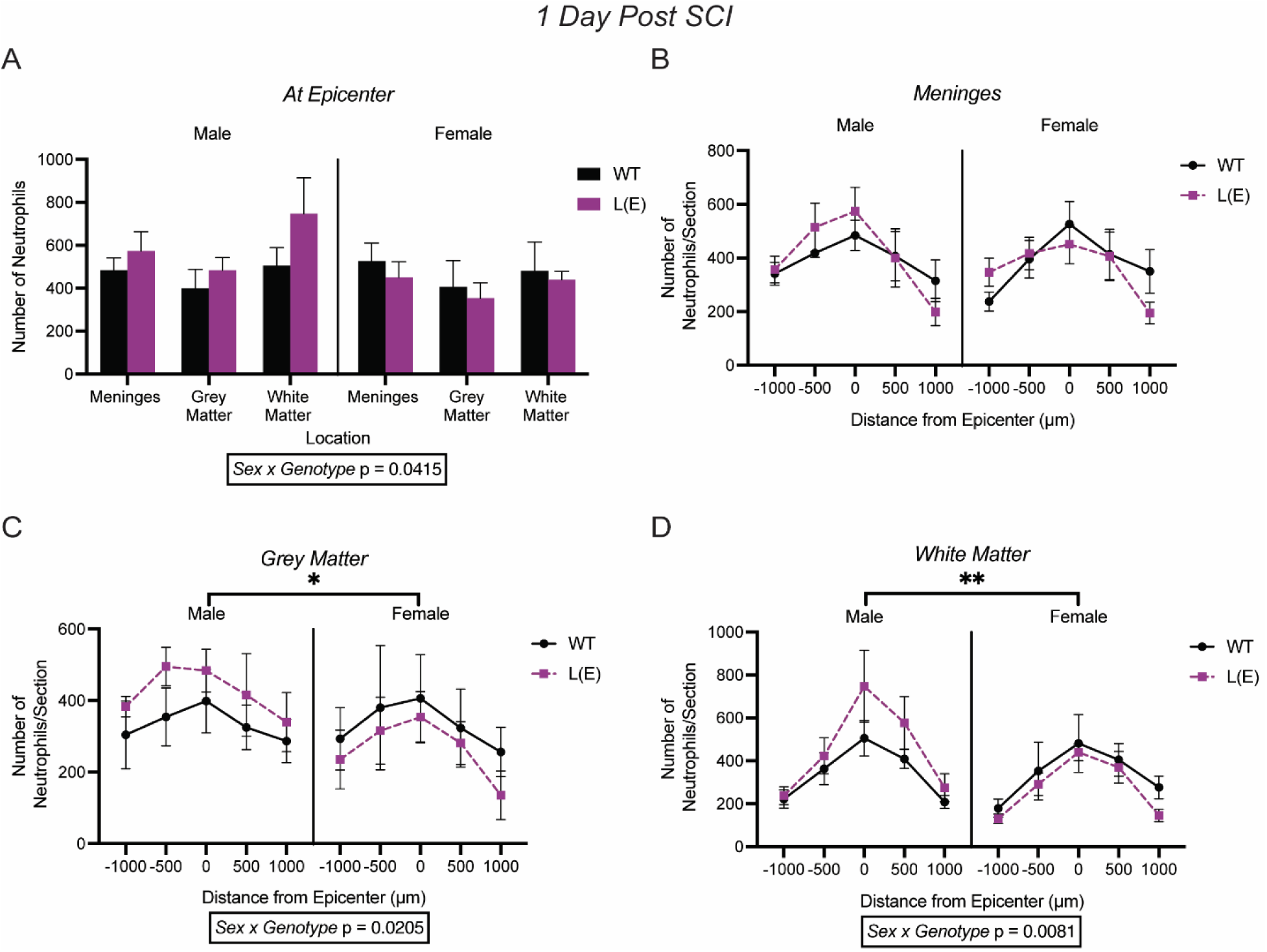
Neutrophil accumulation in the meninges, white matter and grey matter at 1 day after SCI. Neutrophil accumulation at the injury epicenter in the grey matter, white matter, and meninges of the spinal cord. Neutrophil accumulation throughout the injury in the **(B)** meninges, **(C)** grey matter **(D)** white matter. 3-way ANOVA, n = 5-7/sex/genotype, data is shown as mean ± SEM. * p<0.05, ** p<0.01,

**Supplemental Figure 2.**
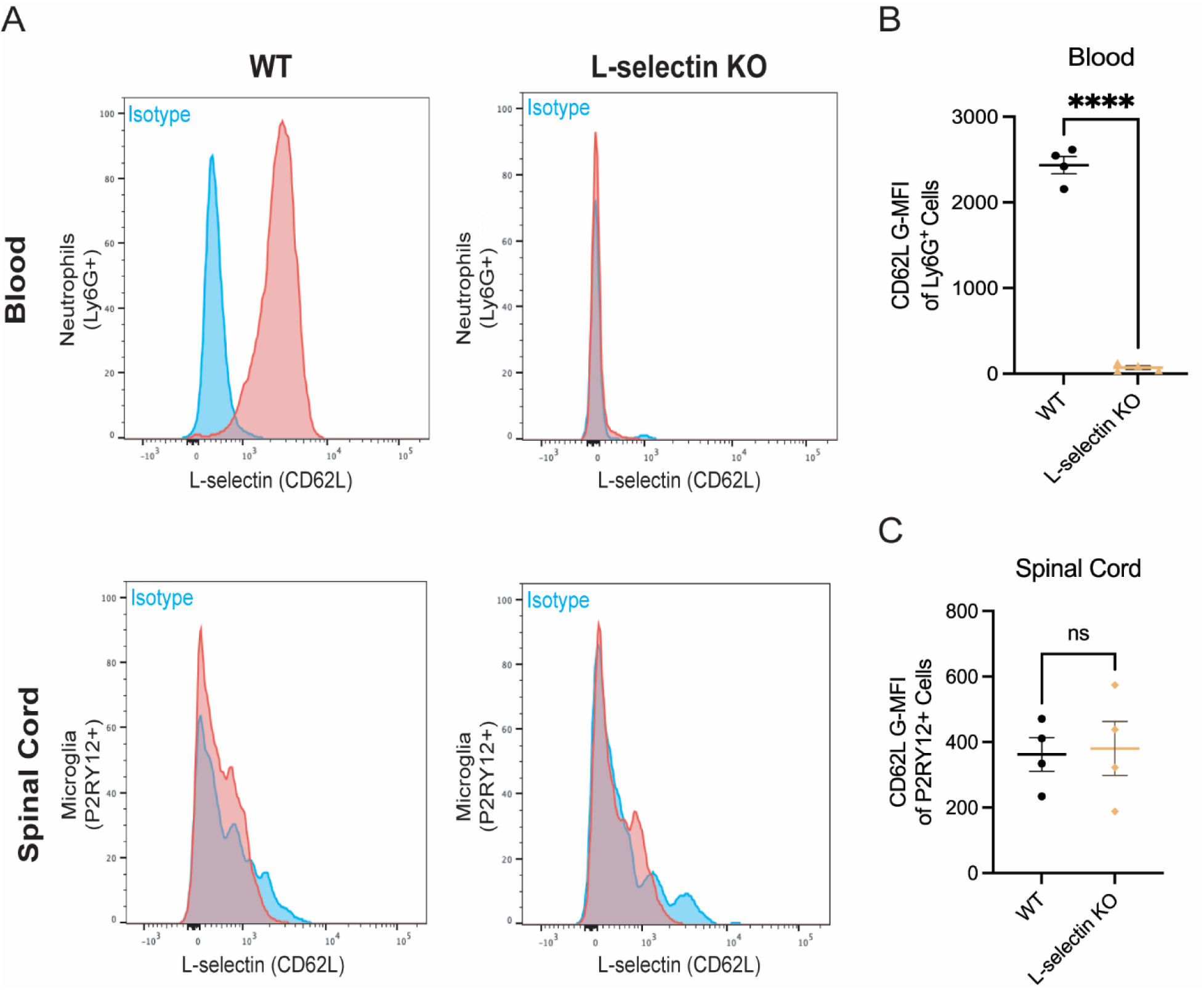
Microglia do not express L-selectin. To determine if microglia express L-selectin, we collected the blood and spinal cord of uninjured male and female WT mice (C57Bl/6N). We also utilized L-selectin KO mice as a control. **(A)** Representative flow cytometry histograms of L-selectin surface expression from the blood and spinal cord of WT and L-selectin KO neutrophils (Ly6G^+^) and microglia (P2RY12^+^). **(B)** In the blood of uninjured mice, neutrophils from WT mice express L-selectin and as expected, neutrophils from L-selectin KO mice do not. **(C)** In the spinal cord of uninjured mice, microglia from WT and L-selectin KO mice have no difference in the G-MFI of L-selectin, indicating that microglia do not express L-selectin. n=4/genotype, data are shown as mean ± SEM,**** p<0.0001

**Supplemental Figure 3.**
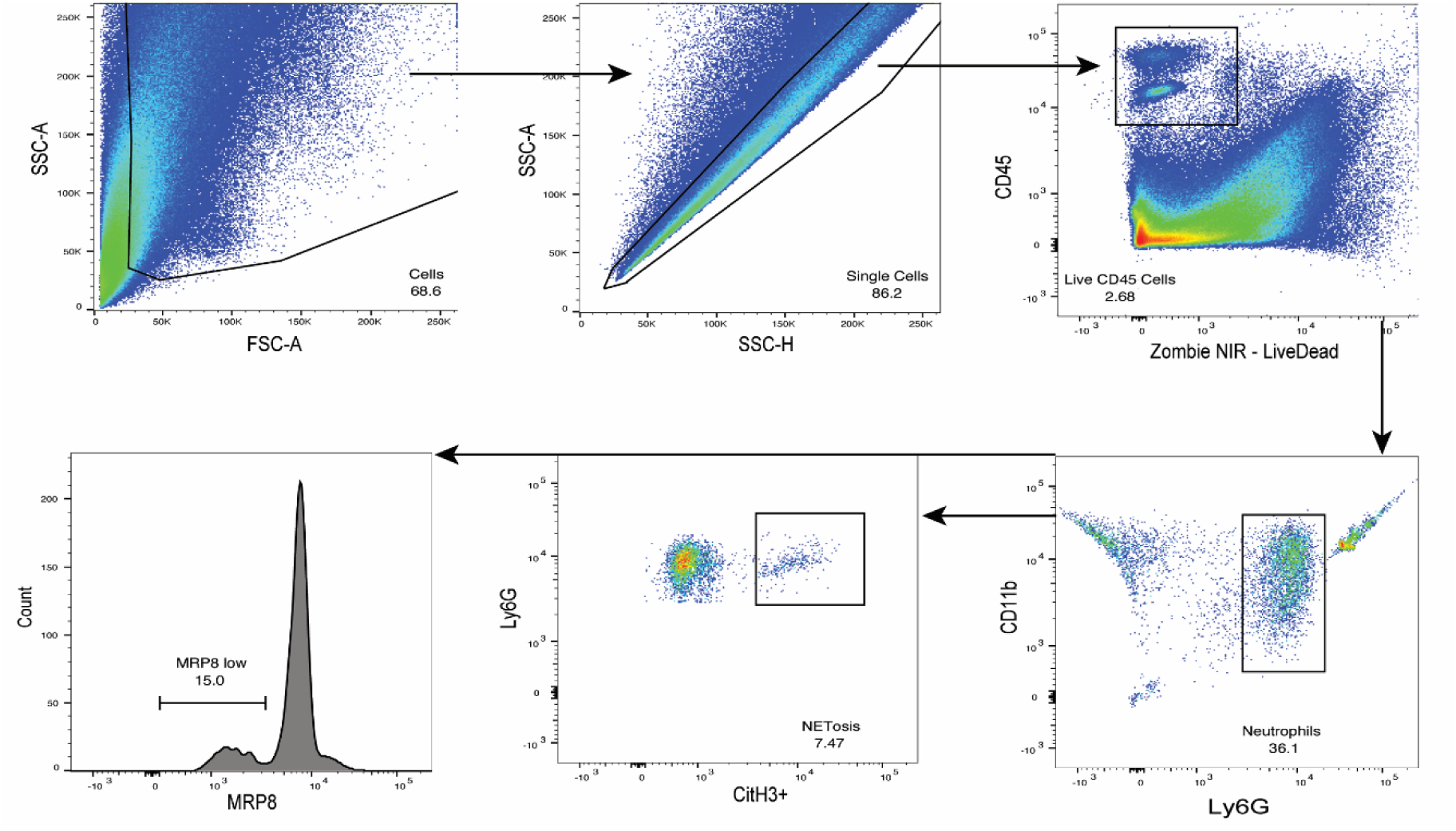
Flow cytometry gating for intracellular staining of neutrophil effector functions at 1 day post injury. Representative images of the gating strategy used to determine neutrophil degranulation (MRP8^low^) and NETosis (CitH3^+^) in the injured spinal cord.

**Supplemental Figure 4.**
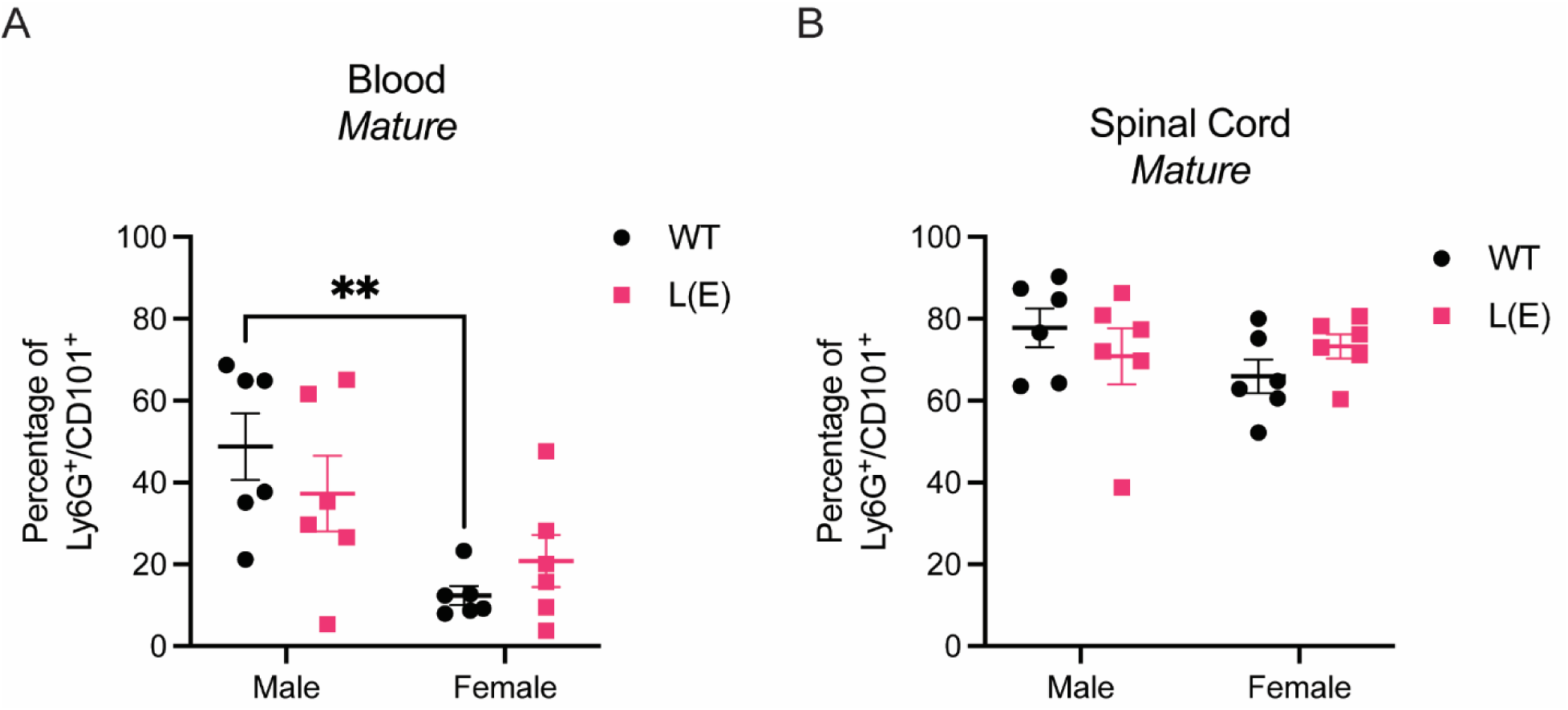
Neutrophil maturiation profiles in the blood and spinal cord 1 day post injury. **(A)** There is no differences in the mature (CD101^+^) neutrophil populations of WT and L(E) mice. Male mice have more mature neutrophils relative to female mice in the blood at 1 day post injury (p_sex_= 0.0012). **(B)** No difference in the percentage of mature instraspinal neutrophil was observed between WT and L(E) mice at 1 day post injury. 2-way ANOVA, n = 6/sex/genotype, data are shown as mean ± SEM. **p<0.01

**Supplemental Figure 5.**
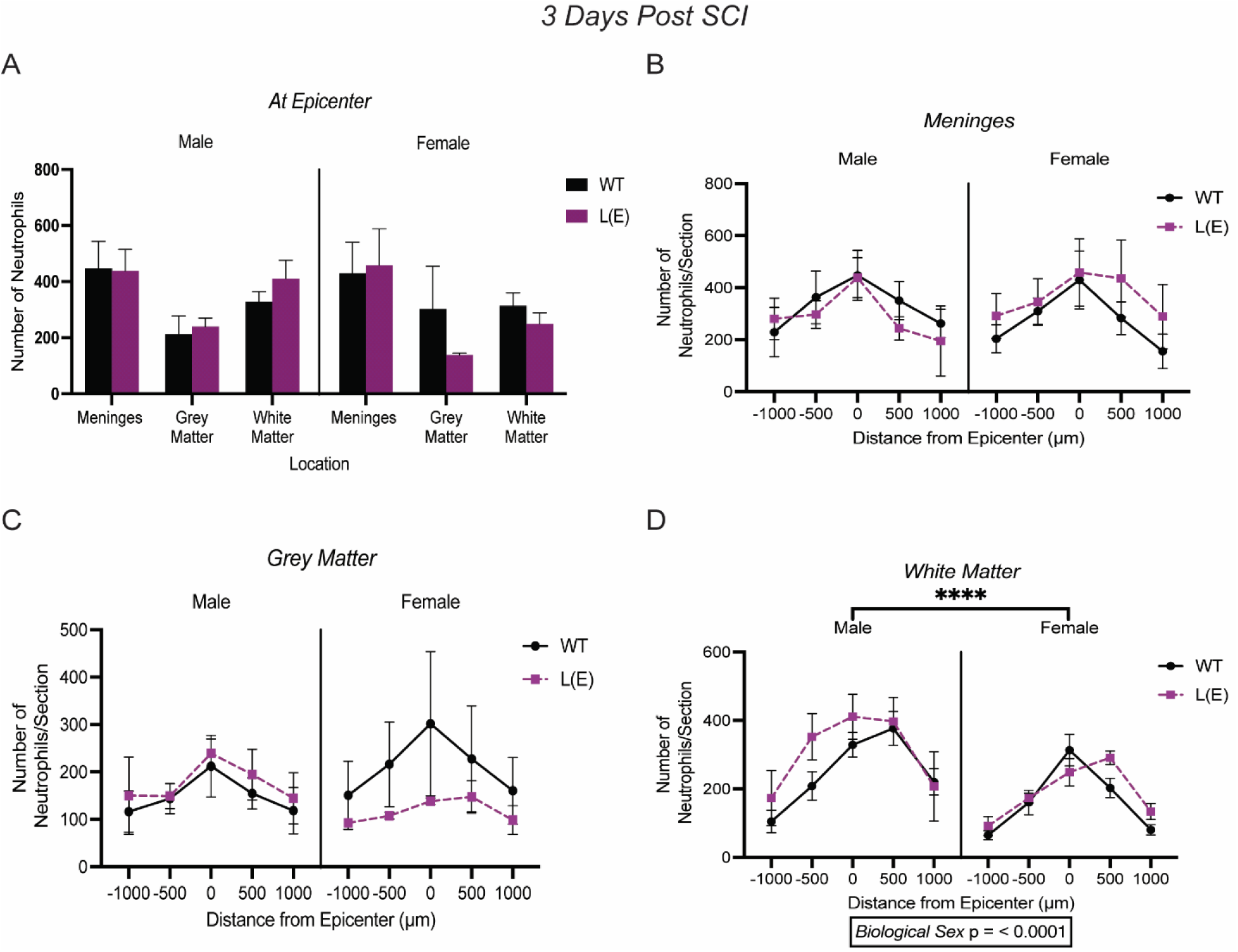
Neutrophil accumulation in the meninges, white matter and grey matter at 3 days post injury. Neutrophil accumulation at the injury epicenter in the grey matter, white matter, and meninges. Neutrophil accumulation throughout the injury in the **(B)** meninges, **(C)** grey matter **(D)** white matter. 3-way ANOVA, n = 3 – 6/sex/genotype, data are shown as mean ± SEM. ****p<0.0001.

**Supplemental Figure 6.**
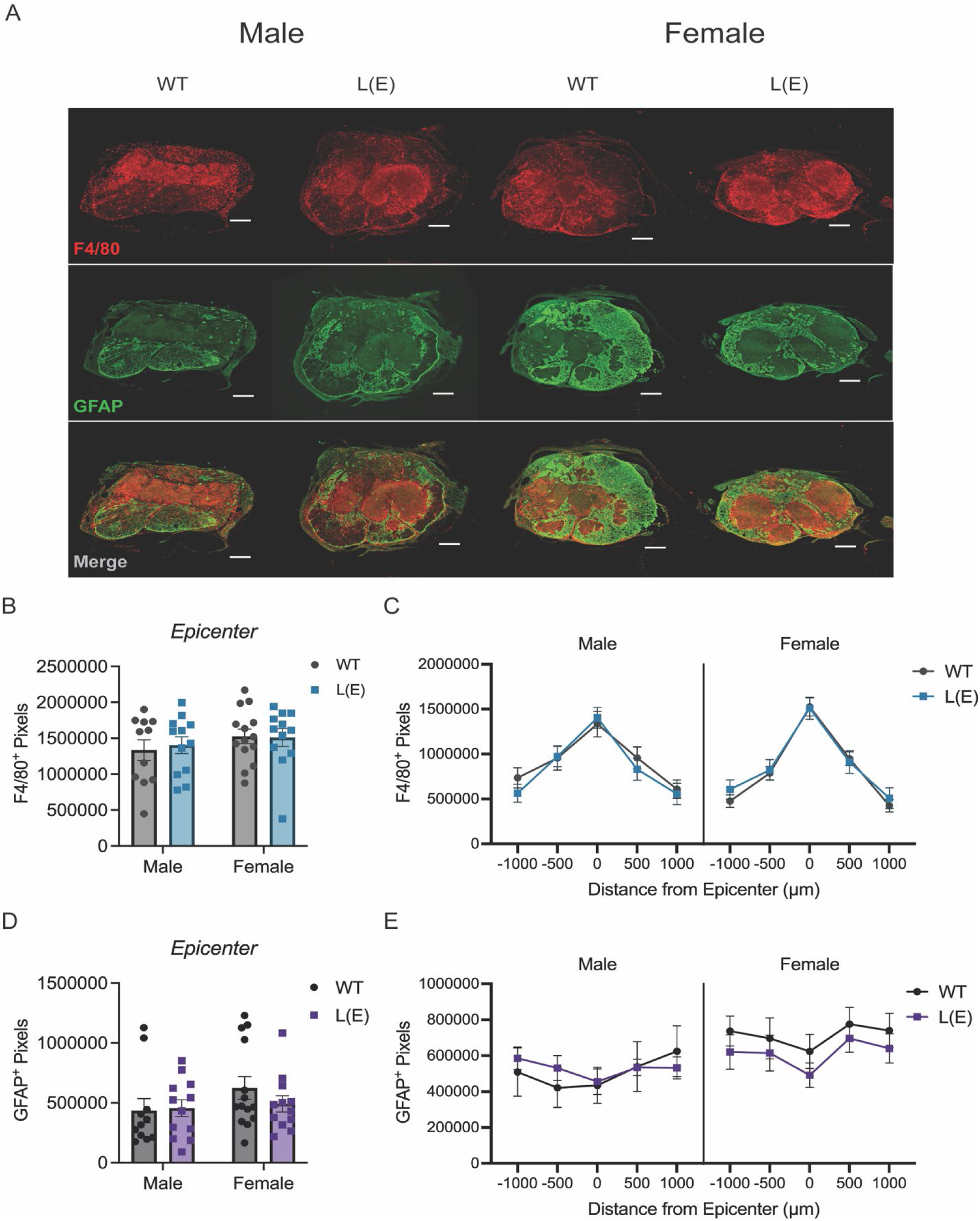
L-selectin shedding does not alter macrophage accumulation or gliosis at 35 days post injury. **(A)** Representative images of macrophages (F4/80) and astrocytes (GFAP) at the lesion epicenter of WT and L(E) male and female mice (scale bar 200μm, 10X) **(B)** Macrophage accumulation (F4/80^+^ pixels) at the lesion epicenter. **(C)** Macrophage accumulation (F4/80^+^ pixels) throughout the injury **(D)** Astrocyte accumulation (GFAP^+^ pixels) at the lesion epicenter. **(E)** Astrocyte accumulation (GFAP^+^ pixels) throughout the injury. n = 11 – 14/sex/genotype, data are shown as mean ± SEM.

## References

1. Alizadeh, A., Dyck, S.M., and Karimi-Abdolrezaee, S. (2019). Traumatic Spinal Cord Injury: An Overview of Pathophysiology, Models and Acute Injury Mechanisms. Front. Neurol. 10. 10.3389/fneur.2019.00282.

2. Pittman, K., and Kubes, P. (2013). Damage-Associated Molecular Patterns Control Neutrophil Recruitment. J. Innate Immun. 5, 315–323. 10.1159/000347132.

3. Scotland, R.S., Stables, M.J., Madalli, S., Watson, P., and Gilroy, D.W. (2011). Sex differences in resident immune cell phenotype underlie more efficient acute inflammatory responses in female mice. Blood 118, 5918–5927. 10.1182/blood-2011-03-340281.

4. Pace, S., Rossi, A., Krauth, V., Dehm, F., Troisi, F., Bilancia, R., Weinigel, C., Rummler, S., Werz, O., and Sautebin, L. (2017). Sex differences in prostaglandin biosynthesis in neutrophils during acute inflammation. Sci. Rep. 7, 3759. 10.1038/s41598-017-03696-8.

5. Gupta, S., Nakabo, S., Blanco, L.P., O’Neil, L.J., Wigerblad, G., Goel, R.R., Mistry, P., Jiang, K., Carmona-Rivera, C., Chan, D.W., et al. (2020). Sex differences in neutrophil biology modulate response to type I interferons and immunometabolism. Proc. Natl. Acad. Sci. U. S. A. 117, 16481–16491. 10.1073/pnas.2003603117.

6. Berghöfer, B., Frommer, T., Haley, G., Fink, L., Bein, G., and Hackstein, H. (2006). TLR7 ligands induce higher IFN-alpha production in females. J. Immunol. Baltim. Md 1950 *177*, 2088–2096. 10.4049/jimmunol.177.4.2088.

7. Beck, K.D., Nguyen, H.X., Galvan, M.D., Salazar, D.L., Woodruff, T.M., and Anderson, A.J. (2010). Quantitative analysis of cellular inflammation after traumatic spinal cord injury: evidence for a multiphasic inflammatory response in the acute to chronic environment. Brain 133, 433–447. 10.1093/brain/awp322.

8. Stirling, D.P., and Yong, V.W. (2008). Dynamics of the inflammatory response after murine spinal cord injury revealed by flow cytometry. J. Neurosci. Res. 86, 1944–1958. 10.1002/jnr.21659.

9. Okada, S. (2016). The pathophysiological role of acute inflammation after spinal cord injury. Inflamm. Regen. 36, 20. 10.1186/s41232-016-0026-1.

10. Neirinckx, V., Coste, C., Franzen, R., Gothot, A., Rogister, B., and Wislet, S. (2014). Neutrophil contribution to spinal cord injury and repair. J. Neuroinflammation 11, 150. 10.1186/s12974-014-0150-2.

11. Donnelly, D.J., and Popovich, P.G. (2008). Inflammation and its role in neuroprotection, axonal regeneration and functional recovery after spinal cord injury. Exp. Neurol. 209, 378–388. 10.1016/j.expneurol.2007.06.009.

12. Kumar, H., Choi, H., Jo, M.-J., Joshi, H.P., Muttigi, M., Bonanomi, D., Kim, S.B., Ban, E., Kim, A., Lee, S.-H., et al. (2018). Neutrophil elastase inhibition effectively rescued angiopoietin-1 decrease and inhibits glial scar after spinal cord injury. Acta Neuropathol. Commun. 6, 73. 10.1186/s40478-018-0576-3.

13. Kubota, K., Saiwai, H., Kumamaru, H., Maeda, T., Ohkawa, Y., Aratani, Y., Nagano, T., Iwamoto, Y., and Okada, S. (2012). Myeloperoxidase exacerbates secondary injury by generating highly reactive oxygen species and mediating neutrophil recruitment in experimental spinal cord injury. Spine 37, 1363–1369. 10.1097/BRS.0b013e31824b9e77.

14. Michel-Flutot, P., Bourcier, C.H., Emam, L., Gasser, A., Glatigny, S., Vinit, S., and Mansart, A. (2023). Extracellular traps formation following cervical spinal cord injury. Eur. J. Neurosci. 57, 692–704. 10.1111/ejn.15902.

15. Savill, J.S., Wyllie, A.H., Henson, J.E., Walport, M.J., Henson, P.M., and Haslett, C. (1989). Macrophage phagocytosis of aging neutrophils in inflammation. Programmed cell death in the neutrophil leads to its recognition by macrophages. J. Clin. Invest. 83, 865–875. 10.1172/JCI113970.

16. Elks, P.M., van Eeden, F.J., Dixon, G., Wang, X., Reyes-Aldasoro, C.C., Ingham, P.W., Whyte, M.K.B., Walmsley, S.R., and Renshaw, S.A. (2011). Activation of hypoxia-inducible factor-1α (Hif-1α) delays inflammation resolution by reducing neutrophil apoptosis and reverse migration in a zebrafish inflammation model. Blood 118, 712–722. 10.1182/blood-2010-12-324186.

17. Fadok, V.A., McDonald, P.P., Bratton, D.L., and Henson, P.M. (1998). Regulation of macrophage cytokine production by phagocytosis of apoptotic and post-apoptotic cells. Biochem. Soc. Trans. 26, 653–656. 10.1042/bst0260653.

18. Hirano, Y., Aziz, M., and Wang, P. (2016). Role of reverse transendothelial migration of neutrophils in inflammation. Biol. Chem. 397, 497–506. 10.1515/hsz-2015-0309.

19. Al-Mayouf, S.M., Sunker, A., Abdwani, R., Abrawi, S.A., Almurshedi, F., Alhashmi, N., Al Sonbul, A., Sewairi, W., Qari, A., Abdallah, E., et al. (2011). Loss-of-function variant in DNASE1L3 causes a familial form of systemic lupus erythematosus. Nat. Genet. 43, 1186– 1188. 10.1038/ng.975.

20. Mahajan, A., Herrmann, M., and Muñoz, L.E. (2016). Clearance Deficiency and Cell Death Pathways: A Model for the Pathogenesis of SLE. Front. Immunol. 7, 35. 10.3389/fimmu.2016.00035.

21. Hakkim, A., Fürnrohr, B.G., Amann, K., Laube, B., Abed, U.A., Brinkmann, V., Herrmann, M., Voll, R.E., and Zychlinsky, A. (2010). Impairment of neutrophil extracellular trap degradation is associated with lupus nephritis. Proc. Natl. Acad. Sci. U. S. A. 107, 9813– 9818. 10.1073/pnas.0909927107.

22. Triantafyllou, E., Pop, O.T., Possamai, L.A., Wilhelm, A., Liaskou, E., Singanayagam, A., Bernsmeier, C., Khamri, W., Petts, G., Dargue, R., et al. (2018). MerTK expressing hepatic macrophages promote the resolution of inflammation in acute liver failure. Gut 67, 333–347. 10.1136/gutjnl-2016-313615.

23. McCreedy, D.A., Abram, C.L., Hu, Y., Min, S.W., Platt, M.E., Kirchhoff, M.A., Reid, S.K., Jalufka, F.L., and Lowell, C.A. (2021). Spleen tyrosine kinase facilitates neutrophil activation and worsens long-term neurologic deficits after spinal cord injury. J. Neuroinflammation 18, 302. 10.1186/s12974-021-02353-2.

24. Ji, J., Zhong, H., Li, Y., Billiar, T.R., Wilson, M.A., Scott, M.J., and Fan, J. (2024). IRG1/ACOD1 promotes neutrophil reverse migration and alleviates local inflammation. J. Leukoc. Biol., qiae110. 10.1093/jleuko/qiae110.

25. Saiwai, H., Ohkawa, Y., Yamada, H., Kumamaru, H., Harada, A., Okano, H., Yokomizo, T., Iwamoto, Y., and Okada, S. (2010). The LTB4-BLT1 axis mediates neutrophil infiltration and secondary injury in experimental spinal cord injury. Am. J. Pathol. 176, 2352–2366. 10.2353/ajpath.2010.090839.

26. Brennan, F.H., Jogia, T., Gillespie, E.R., Blomster, L.V., Li, X.X., Nowlan, B., Williams, G.M., Jacobson, E., Osborne, G.W., Meunier, F.A., et al. (2019). Complement receptor C3aR1 controls neutrophil mobilization following spinal cord injury through physiological antagonism of CXCR2. JCI Insight 4, 98254. 10.1172/jci.insight.98254.

27. Reid, S.K., Leal-Garcia, M.E., Tran, A.V., Rehtmeyer, N.T., Shirvaikar, I.S., Kirchhoff, M.A., Narvaez, A.O., and McCreedy, D.A. (2025). Recombinant human DNase treatment mitigates extracellular trap mediated damage and improves long-term recovery after spinal cord injury in male mice. Brain. Behav. Immun. 128, 456–468. 10.1016/j.bbi.2025.04.033.

28. Ivetic, A. (2018). A head-to-tail view of L-selectin and its impact on neutrophil behaviour. Cell Tissue Res. 371, 437–453. 10.1007/s00441-017-2774-x.

29. Grailer, J.J., Kodera, M., and Steeber, D.A. (2009). L-selectin: Role in Regulating Homeostasis and Cutaneous Inflammation. J. Dermatol. Sci. 56, 141–147. 10.1016/j.jdermsci.2009.10.001.

30. Tedder, T.F., Steeber, D.A., and Pizcueta, P. (1995). L-selectin-deficient mice have impaired leukocyte recruitment into inflammatory sites. J. Exp. Med. 181, 2259–2264. 10.1084/jem.181.6.2259.

31. Arbonés, M.L., Ord, D.C., Ley, K., Ratech, H., Maynard-Curry, C., Otten, G., Capon, D.J., and Teddert, T.F. (1994). Lymphocyte homing and leukocyte rolling and migration are impaired in L-selectin-deficient mice. Immunity 1, 247–260. 10.1016/1074-7613(94)90076-0.

32. Crockett-Torabi, E., Sulenbarger, B., Smith, C.W., and Fantone, J.C. (1995). Activation of human neutrophils through L-selectin and Mac-1 molecules. J. Immunol. Baltim. Md 1950 *154*, 2291–2302.

33. Laudanna, C., Constantin, G., Baron, P., Scarpini, E., Scarlato, G., Cabrini, G., Dechecchi, C., Rossi, F., Cassatella, M.A., and Berton, G. (1994). Sulfatides trigger increase of cytosolic free calcium and enhanced expression of tumor necrosis factor-alpha and interleukin-8 mRNA in human neutrophils. Evidence for a role of L-selectin as a signaling molecule. J. Biol. Chem. 269, 4021–4026.

34. Cappenberg, A., Margraf, A., Thomas, K., Bardel, B., McCreedy, D.A., Van Marck, V., Mellmann, A., Lowell, C.A., and Zarbock, A. (2019). L-selectin shedding affects bacterial clearance in the lung: a new regulatory pathway for integrin outside-in signaling. Blood 134, 1445–1457. 10.1182/blood.2019000685.

35. Smolen, J.E., Petersen, T.K., Koch, C., O’Keefe, S.J., Hanlon, W.A., Seo, S., Pearson, D., Fossett, M.C., and Simon, S.I. (2000). L-selectin signaling of neutrophil adhesion and degranulation involves p38 mitogen-activated protein kinase. J. Biol. Chem. 275, 15876– 15884. 10.1074/jbc.M906232199.

36. Woodfin, A., Voisin, M.-B., Beyrau, M., Colom, B., Caille, D., Diapouli, F.-M., Nash, G.B., Chavakis, T., Albelda, S.M., Rainger, G.E., et al. (2011). The junctional adhesion molecule JAM-C regulates polarized transendothelial migration of neutrophils in vivo. Nat. Immunol. 12, 761–769. 10.1038/ni.2062.

37. Waddell, T.K., Fialkow, L., Chan, C.K., Kishimoto, T.K., and Downey, G.P. (1994). Potentiation of the oxidative burst of human neutrophils. A signaling role for L-selectin. J. Biol. Chem. 269, 18485–18491. 10.1016/S0021-9258(17)32335-9.

38. Shao, B., Wahrenbrock, M.G., Yao, L., David, T., Coughlin, S.R., Xia, L., Varki, A., and McEver, R.P. (2011). Carcinoma mucins trigger reciprocal activation of platelets and neutrophils in a murine model of Trousseau syndrome. Blood 118, 4015–4023. 10.1182/blood-2011-07-368514.

39. Mohanty, T., Sjögren, J., Kahn, F., Abu-Humaidan, A.H.A., Fisker, N., Assing, K., Mörgelin, M., Bengtsson, A.A., Borregaard, N., and Sørensen, O.E. (2015). A novel mechanism for NETosis provides antimicrobial defense at the oral mucosa. Blood 126, 2128–2137. 10.1182/blood-2015-04-641142.

40. Griffin, J.D., Spertini, O., Ernst, T.J., Belvin, M.P., Levine, H.B., Kanakura, Y., and Tedder, T.F. (1990). Granulocyte-macrophage colony-stimulating factor and other cytokines regulate surface expression of the leukocyte adhesion molecule-1 on human neutrophils, monocytes, and their precursors. J. Immunol. Baltim. Md 1950 *145*, 576–584.

41. Spertini, O., Freedman, A.S., Belvin, M.P., Penta, A.C., Griffin, J.D., and Tedder, T.F. (1991). Regulation of leukocyte adhesion molecule-1 (TQ1, Leu-8) expression and shedding by normal and malignant cells. Leukemia 5, 300–308.

42. Zhao, L.C., Edgar, J.B., and Dailey, M.O. (2001). Characterization of the rapid proteolytic shedding of murine L-selectin. Dev. Immunol. 8, 267–277. 10.1155/2001/91831.

43. Wang, Y., Zhang, A.C., Ni, Z., Herrera, A., and Walcheck, B. (2010). ADAM17 Activity and Other Mechanisms of Soluble L-selectin Production during Death Receptor-Induced Leukocyte Apoptosis. J. Immunol. 184, 4447–4454. 10.4049/jimmunol.0902925.

44. Greenlee-Wacker, M.C. (2016). Clearance of apoptotic neutrophils and resolution of inflammation. Immunol. Rev. 273, 357–370. 10.1111/imr.12453.

45. Strausbaugh, H.J., and Rosen, S.D. (2001). A Potential Role for Annexin 1 as a Physiologic Mediator of Glucocorticoid-Induced L-Selectin Shedding from Myeloid Cells. J. Immunol. 166, 6294–6300. 10.4049/jimmunol.166.10.6294.

46. Buckley, C.D., Ross, E.A., McGettrick, H.M., Osborne, Chloe.E., Haworth, O., Schmutz, C., Stone, P.C.W., Salmon, M., Matharu, N.M., Vohra, R.K., et al. (2006). Identification of a phenotypically and functionally distinct population of long-lived neutrophils in a model of reverse endothelial migration. J. Leukoc. Biol. 79, 303–311. 10.1189/jlb.0905496.

47. McCreedy, D.A., Lee, S., Sontag, C.J., Weinstein, P., Olivas, A.D., Martinez, A.F., Fandel, T.M., Trivedi, A., Lowell, C.A., Rosen, S.D., et al. (2018). Early Targeting of L-Selectin on Leukocytes Promotes Recovery after Spinal Cord Injury, Implicating Novel Mechanisms of Pathogenesis. eNeuro 5, ENEURO.0101-18.2018. 10.1523/ENEURO.0101-18.2018.

48. Venturi, G.M., Tu, L., Kadono, T., Khan, A.I., Fujimoto, Y., Oshel, P., Bock, C.B., Miller, A.S., Albrecht, R.M., Kubes, P., et al. (2003). Leukocyte migration is regulated by L-selectin endoproteolytic release. Immunity 19, 713–724. 10.1016/s1074-7613(03)00295-4.

49. Basso, D.M., Fisher, L.C., Anderson, A.J., Jakeman, L.B., Mctigue, D.M., and Popovich, P.G. (2006). Basso Mouse Scale for Locomotion Detects Differences in Recovery after Spinal Cord Injury in Five Common Mouse Strains. J. Neurotrauma 23, 635–659. 10.1089/neu.2006.23.635.

50. Schleiffenbaum, B., Spertini, O., and Tedder, T.F. (1992). Soluble L-selectin is present in human plasma at high levels and retains functional activity. J. Cell Biol. 119, 229–238. 10.1083/jcb.119.1.229.

51. Spertini, O., Schleiffenbaum, B., White-Owen, C., Ruiz, P., and Tedder, T.F. (1992). ELISA for quantitation of L-selectin shed from leukocytes in vivo. J. Immunol. Methods 156, 115–123. 10.1016/0022-1759(92)90017-n.

52. Tu, L., Poe, J.C., Kadono, T., Venturi, G.M., Bullard, D.C., Tedder, T.F., and Steeber, D.A. (2002). A functional role for circulating mouse L-selectin in regulating leukocyte/endothelial cell interactions in vivo. J. Immunol. Baltim. Md 1950 *169*, 2034–2043. 10.4049/jimmunol.169.4.2034.

53. Adrover, J.M., Del Fresno, C., Crainiciuc, G., Cuartero, M.I., Casanova-Acebes, M., Weiss, L.A., Huerga-Encabo, H., Silvestre-Roig, C., Rossaint, J., Cossío, I., et al. (2019). A Neutrophil Timer Coordinates Immune Defense and Vascular Protection. Immunity 50, 390–402.e10. 10.1016/j.immuni.2019.01.002.

54. Aubé, B., Lévesque, S.A., Paré, A., Chamma, É., Kébir, H., Gorina, R., Lécuyer, M.-A., Alvarez, J.I., De Koninck, Y., Engelhardt, B., et al. (2014). Neutrophils Mediate Blood– Spinal Cord Barrier Disruption in Demyelinating Neuroinflammatory Diseases. J. Immunol. 193, 2438–2454. 10.4049/jimmunol.1400401.

55. Lee, J.Y., Choi, H.Y., Ahn, H.-J., Ju, B.G., and Yune, T.Y. (2014). Matrix Metalloproteinase-3 Promotes Early Blood–Spinal Cord Barrier Disruption and Hemorrhage and Impairs Long-Term Neurological Recovery after Spinal Cord Injury. Am. J. Pathol. 184, 2985–3000. 10.1016/j.ajpath.2014.07.016.

56. Kuhn, P.L., and Wrathall, J.R. (1998). A mouse model of graded contusive spinal cord injury. J. Neurotrauma 15, 125–140. 10.1089/neu.1998.15.125.

57. Jeon, S.-B., Yoon, H.J., Park, S.-H., Kim, I.-H., and Park, E.J. (2008). Sulfatide, A Major Lipid Component of Myelin Sheath, Activates Inflammatory Responses As an Endogenous Stimulator in Brain-Resident Immune Cells1. J. Immunol. 181, 8077–8087. 10.4049/jimmunol.181.11.8077.

58. Haage, V., Semtner, M., Vidal, R.O., Hernandez, D.P., Pong, W.W., Chen, Z., Hambardzumyan, D., Magrini, V., Ly, A., Walker, J., et al. (2019). Comprehensive gene expression meta-analysis identifies signature genes that distinguish microglia from peripheral monocytes/macrophages in health and glioma. Acta Neuropathol. Commun. 7, 20. 10.1186/s40478-019-0665-y.

59. Hu, X., Liou, A.K.F., Leak, R.K., Xu, M., An, C., Suenaga, J., Shi, Y., Gao, Y., Zheng, P., and Chen, J. (2014). Neurobiology of microglial action in CNS injuries: receptor-mediated signaling mechanisms and functional roles. Prog. Neurobiol. 0, 60–84. 10.1016/j.pneurobio.2014.06.002.

60. Simon, S.I., Burns, A.R., Taylor, A.D., Gopalan, P.K., Lynam, E.B., Sklar, L.A., and Smith, C.W. (1995). L-selectin (CD62L) cross-linking signals neutrophil adhesive functions via the Mac-1 (CD11b/CD18) beta 2-integrin. J. Immunol. Baltim. Md 1950 155, 1502–1514.

61. Abraham, E., Carmody, A., Shenkar, R., and Arcaroli, J. (2000). Neutrophils as early immunologic effectors in hemorrhage- or endotoxemia-induced acute lung injury. Am. J. Physiol.-Lung Cell. Mol. Physiol. 279, L1137–L1145. 10.1152/ajplung.2000.279.6.L1137.

